# Lipidated poly(amino acid) nanostructures as versatile therapeutic delivery vehicles

**DOI:** 10.1101/2020.04.07.004333

**Authors:** Josiah D. Smith, Leah N. Cardwell, David Porciani, Andrew J. Greenwald, Aiden C. Ellis, Megan C. Schulte, Xiaofei Wang, Evan T. Schoenherr, Gracen F. Seim, Joe E. Anderson, Julie A. Nguyen, Rama R. Tata, Margaret J. Lange, Donald H. Burke, Mark A. Daniels, Bret D. Ulery

## Abstract

Poly(amino acid)s are a diverse and capable class of polymers with significant potential for utilization in a wide variety of drug delivery applications. A sub-class of these biomaterials known as lipidated poly(amino acid)s (LPAAs) are amphiphiles composed of both hydrophobic and hydrophilic domains yielding interesting physical properties. In this article, we describe our efforts in developing a novel class of lysine and valine containing LPAAs synthesized via hexadecylamine initiated N-carboxyanhydride ring-opening polymerization (NCA-ROP). These highly hydrophobic LPAAs were found capable of undergoing hydrophobically-driven self-assembly into small nanostructures as well as being forced into larger nanostructures using a novel dump-and-stir nanoprecipitation process. This process yielded fine control over resulting nanoparticle size and cargo entrapment. Furthermore, cell-targeting DNA aptamer modification of doxorubicin-loaded LPAA nanoparticles induced significant death of co-incubated Non-Hodgkin Lymphoma cells providing exciting evidence of the therapeutic potential of this novel biomaterials-based delivery device.

## Introduction

Non-Hodgkin Lymphoma (NHL) is a common, deadly cancer with an estimated 74,000 new cases and 20,000 deaths in the United States in 2018^1^. Approximately 60% of NHLs are aggressive subtypes likely to spread to several lymph nodes, peripheral tissues, and other organs causing harder to treat and more deadly disease^2^. With only a 71% 5-year survival rate, there exists a need for novel therapeutics to combat this cancer^3^. Addressing the diffuse and metastatic nature of the most common hematological cancer subtype, B cell NHL, the clinical standard therapy is a lymphocyte specific monoclonal antibody concurrent with combination chemotherapy, known as R-CHOP (rituximab, cyclophosphamide, doxorubicin, vincristine and prednisolone)^4^. While somewhat therapeutically effective, R-CHOP is associated with considerable side-effects including chemotherapeutic toxicity, permanent fatigue, and potential lifelong immunodeficiency^5–7^.

A strategy to address the off-target toxicity of chemotherapeutics is cell-targeted delivery. For example, the antibody-drug conjugate composed of Inotuzumab (antibody) and ozogamicin (drug), is used for the targeted delivery of a calicheamicin, a highly toxic agent to all cell types, directly to CD22, a B cell surface marker^8^. This therapy was FDA approved in 2017 for the treatment of leukemia and is currently undergoing clinical trials for NHL^9^. Due to the complex and costly nature of antibody-drug conjugates, a simpler targeting moiety would be valuable^10^. An alternative to using antibodies for cell-targeted chemotherapeutic delivery is employing aptamers, which are nucleic acid oligomers capable of selectively binding target biological sites^11^. Aptamers are attractive for targeted drug delivery due to their favorable toxicity profiles and lower immunogenicity compared with antibodies^12^. Aptamers have been identified that bind CD20 (*e.g*., DNA aptamer ACDA), the same receptor targeted by the antibody drug rituximab, several B cell lymphoma and leukemia cells lines (*e.g*., DNA aptamer C10.36), and transferrin receptor (*e.g*., RNA aptamer WAZ)^13–16^.

A method to improve the delivery capacity of an aptamer or antibody by attachment to the surface of a drug-loaded nanoparticle. An additional benefit of this approach is that nanoparticle-based drug entrapment has been shown to reduce chemotherapeutic side-effects, as evidenced by the use of liposomes to deliver doxorubicin for the treatment of breast cancer^17^. In choosing which nanoparticle system to use, material composition must be suitable for the intended application, with degradation mechanism and kinetics appropriate for the desired biological destination. While many materials have been studied, polymers are attractive as they can be made from a variety of chemistries with varying degradation mechanisms including commonly used ones like poly(lactide-co-glyocolide), hyaluronan, poly(dopamine), and poly(aspartic acid)^18–21^.

Poly(amino acid)s are an interesting and unique class of polymer that have been explored for their drug delivery potential. First developed in the 1950s, N-carboxyanhydride (NCA)-based poly(amino acid)s are producible at large scales, have highly controlled molecular weights, and can be readily made from a diverse set of functional groups^22–25^. Also, poly(amino acid)s have been previously used to form nanoparticles for drug entrapment^26, 27^. A subtype of this polymer class are lipidated poly(amino acid)s (LPAAs) which are characterized as possessing a hydrophilic peptide linked to a hydrophobic tail and are able to undergo hydrophobically-driven self-assembled into nanostructures^28–31^ similar to a more commonly studied class of materials in peptide amphiphiles.

In this article, we describe our design of novel, hydrophobic NCA-derived LPAAs comprised of lysine and valine. The lipid hexadecylamine was used as initiator, resulting in a set of poly(K_a_V_b_)C_16_ polymers with varying lysine/valine ratios (*i.e*., 30/30, 15/45, 6/53, 3/57) with precise molecular weights and an average of 60 amino acids per polymer. These LPAAs were found to selfassemble into micelles (LPAAMs) in phosphate buffered saline (PBS). To best entrap drug payloads within their nanostructures, a novel nanoprecipitation system was developed. LPAA and drugs were co-dissolved in dimethyl formamide (DMF) and precipitated in a mixture of octanol and pentane. The resulting lipidated poly(amino acid) nanoparticles (LPAANPs) were capable of successfully entrapping doxorubicin (Dox), peptide amphiphile (PA), or hydrophilic peptide (P), each with high efficiency. Cell safety studies were completed to evaluate the compatibility of LPAA nanostructures with murine fibroblasts, for which limited toxicity was observed with LPAAMs (only at high doses – 500 and 1000 μg/mL) and no toxicity was seen with LPAANPs. Finally, a Dox-loaded LPAANP therapeutic delivery vehicle was surface-modified to display aptamer C10.36 for cell specific drug targeting. These modified nanostructures were found to possess enhanced cellular association with and cytotoxicity against a human NHL cell line (*i.e*., Ramos cells). As LPAANPs can readily entrap other therapeutics and/or be surface modified with different targeting moieties, this novel drug delivery device can serve as a platform technology for a variety of biomedical applications.

## Methods

### Polymer Synthesis

#### N-Carboxyanhydride (NCA) Synthesis

All reagents and solvents were purchased from Sigma-Aldrich unless specified otherwise. H_2_N-lysine(carboxybenzyl)-OH (lysine(Cbz)) and H_2_N-valine-OH (valine) were dried for 48 hours under hard vacuum (Welch 8912 direct drive pump) prior to use. Tetrahydrofuran (THF) was dried with molecular sieves for at least 24 hours prior to use. Dry starting material (4 g / 14 mmol lysine(Cbz) or 4 g / 34 mmol valine) was added to a 250 mL round bottom flask (RBF) along with 20 mL of THF after which the reaction mixture was warmed to 55°C in an oil bath under constant magnetic stirring. A solution of triphosgene at 0.5 mmol triphosgene per mmol amino acid was produced by dissolving triphosgene in 10 mL of THF and added slowly to the RBF. Additional THF (25 mL) was added to rinse the walls of the RBF as each amino acid used was initially minimally soluble in THF. The RBF was capped to allow for the gentle condensing reflux of THF to rinse the RBF walls. The Lysine(Cbz) reaction solution becomes transparent after 45 minutes and the reaction was stopped after 60 minutes. Valine reaction solution becomes transparent by 105 minutes and the reaction was stopped after 120 minutes. Each RBF was cooled to room temperature and solvent removed via rotary evaporation (Büchi R-205) until ^~^ 5 mL was left. Hexane (100 mL) was added and crude N-carboxyanhydride (NCA) of each amino acid precipitated at room temperature within 10 minutes. The hexane solution was decanted and each crude NCA was washed with an additional 50 mL of hexane three times to remove residual triphosgene and THF. Residual hexane was removed from each RBF by rotary evaporation followed by drying under high vacuum (Welch 8912 direct drive pump) for 30 minutes. A scheme for each reaction and corresponding yields are shown in **Figure S1**.

#### Purification

Ethyl acetate was dried under anhydrous sodium sulfate for 10 minutes prior to use. Silica was dried under vacuum (Büchi V-700 diaphragm pump) for 48 hours prior to use. Crude NCAs were dissolved in a 250 mL RBF in ethyl acetate at ^~^ 500 mg/mL with gentle heating to 55°C. Dry, powdered silica (^~^ 50 mL) was added and residual ethyl acetate removed by rotary evaporation. The target molecule was isolated by purification on a silica gel column containing a total loose volume of 75 mL dry silica with a mixture of ethyl acetate and hexane as the mobile phase ^32^. NCAs eluted at ^~^ 50 - 90% ethyl acetate in hexane dependent on the NCA species. Each pure NCA was dried via rotary evaporation and under hard vacuum (Welch 8912 direct drive pump) for 1 hour. NCAs were kept on ice and used for polymerization within 1 hour of their synthesis.

#### N-Carboxyanhydride Ring Opening Polymerization (NCA-ROP)

NCAs were dissolved in fresh dimethylformamide (DMF) at 250 mg/mL. NCAs were mixed in fixed molar ratios according to the desired final ratio for the resulting polymer (*i.e*., lysine/valine ratios of 30/30, 15/45, 6/54, and 3/57, resulting in LPAAs with molar percentages of lysine compared to total amino acid content of 50%, 25%, 10%, and 5%). NCA mixtures were then added dropwise with constant stirring to a solution containing hexadecylamine initiator at 25 mg/mL in 5 mL of a 1:1 mixture of DMF and chloroform (CHCl_3_) in a 25 mL RBF on ice. The molar ratio of hexadecylamine to total NCA added was held at 1:60. Each RBF was sealed with a rubber septa with a small opening in each cap for the release of carbon dioxide that evolves during NCA-ROP. Each reaction proceeded overnight under constant stirring and slowly warmed to room temperature. The reaction and related information for this process are shown in **Scheme S1**. DMF and CHCl_3_ were removed via rotary evaporation followed by the addition of 5 mL of dichloromethane (DCM) and further rotary evaporation to assist in the removal of residual DMF. Each protected LPAA was dried under hard vacuum overnight prior to analysis and deprotection.

#### Deprotection

The LPAA lysine carboxybenzyl (Cbz) protection group was removed by treatment with hydrobromic acid (HBr)^33^. Protected LPAA (250 mg) was dissolved in 2.5 mL of trifluoroacetic acid (TFA) at 100 mg/mL in a 25 mL RBF under continuous magnetic stirring. HBr in acetic acid (0.5 mL of a 33% wt/v solution) was added to facilitate the reaction and after 1.5 hours 0.5 mL of HBr in ddH_2_O (48% wt/v solution in distilled, deionized water) was included. Finally, a second 0.5 mL of HBr in ddH_2_O was provided after 3 hours of total deprotection time. The reaction and related information for this process are shown in **Scheme S2**. After 6 hours of deprotection, the reaction was terminated by the removal of solvent via rotary evaporation until ^~^ 1 mL was left. The deprotected LPAAs were immediately precipitated via the addition of 15 mL of diethyl ether. Precipitated LPAA was rinsed with an additional 100 mL of diethyl ether using vacuum filtration. LPAA was then dissolved at 25 mg/mL in ddH_2_O and neutralized to pH 7 via the dropwise addition of 1 M sodium hydroxide as monitored with pH paper. Neutralized polymer solution was frozen and lyophilized using a Labconco FreeZone 4.5.

### LPAA Evaluation

#### Nuclear Magnetic Resonance - Diffusion Ordered Spectroscopy (NMR DOSY)

Protected LPAA was dissolved in 2.5% deuterated TFA (d-TFA) in deuterated chloroform (CDCl_3_) at 1 mg/mL. A portion of this solution (250 μL) was evaluated in a 3 mm NMR tube at 25 °C using an NMR spectrometer (Bruker Avance III 600 MHz) equipped with a cryoprobe. Adequate signal attenuation was assured for each sample via repeated analyses of the signal. Specifically, signal strength was compared for 5% and 95% gradient strength to assure at least 95% of signal attenuation was obtained. Both the diffusion time and gradient length were adjusted as necessary. For a single DOSY acquisition, sixteen proton (^1^H) NMR spectra were taken and compiled into a 2-D DOSY plot using Bruker NMR Software. LPAA diffusion coefficients were determined by plotting diffusivity against chemical shift with LPAA identified by the expected ppm of functional groups (including Cbz and hexadecylamine). Known molecular weights of polystyrene (Agilent GPC/SEC Standard Calibration Kit) were also evaluated by NMR DOSY using the same procedure. A standard curve of diffusivity versus known molecular weights of polystyrene to solve for the molecular weights for LPAAs from experimentally determine diffusion coefficients. LPAA NMR DOSY was completed for three independently produced batches of each LPAA to determine an approximate average molecular weight.

#### LPAA Lysine/Valine Ratio Determination

Protected LPAAs were evaluated to assess their relative lysine and valine content. Protected LPAAs were dissolved in deuterated dimethyl sulfoxide (DMSO-d6) at 10 mg/mL. A portion of this solution (250 μL) in a 3 mm NMR tube at 25 °C was used to generate ^1^H NMR spectrum employing the aforementioned 600 MHz NMR. Relative lysine(Cbz) content was calculated as half of the integration value of the peak at 4.2 ppm, corresponding to the 2 aliphatic hydrogens on the carbon adjacent to the Cbz ring. Relative valine content was estimated as the integration value of the peak at 3.0 ppm, corresponding to the singly hydrogen on the ⍰-carbon associated with the valine backbone. Lysine/valine ratios were assessed for each LPAA from three independently produced batches.

#### LPAA Deprotection Confirmation

Deprotected LPAAs were dissolved in DMSO-d6 at 10 mg/mL and evaluated similarly to protected LPAAs via ^1^H NMR. The hydrogen peaks from the Cbz ring at 7.2 ppm and -CH_2_- peak adjacent to the Cbz ring at 5 ppm were assessed to confirm deprotection. NMR spectra was taken for each LPAA to assure deprotection.

#### Critical Micelle Concentration (CMC)

The CMC, or minimum concentration at which LPAAs self-assemble to form nanoparticles, was assessed using established protocols^34, 35^. LPAAs were dissolved in pH 7.4 phosphate buffered saline (PBS) at 100 μg/mL and serially diluted in 1 μM 1,6-Diphenyl-1,3,5-hexatriene (DPH) in PBS. Each sample was incubated at room temperature for 1 hour. DPH fluorescence was measured using a BioTek Cytation 5 plate reader with an excitation/emission of 350/428 nm. DPH has been shown to stack within hydrophobic pockets, such as the core of a micelle, dramatically increasing in fluorescence. DPH fluorescence was used as a proxy for micelle formation. When plotted as fluorescence against logarithmic concentration, the inflection point in fluorescence was taken as the CMC value. CMCs were taken in triplicate from three independently produced LPAAs of each lysine/valine ratio.

#### Transmission Electron Microscopy (TEM)

Micelle morphology was assessed via TEM. Each sample (5 μL at 1 mg/mL in PBS) was incubated on a carbon-coated copper grid (Pelco 200 mesh) for 5 minutes, followed by wicking with filter paper and treatment with 5 μL of NanoW negative stain. Following 5 minutes of stain treatment, samples were wicked until dry. Both wicking steps were completed using a deep staining method, where the filter paper was applied to the bottom of the grid and sample/stain solution drawn through the grid. Samples were then evaluated via JEOL JEM-1400 TEM with micrographs captured at 120 keV. TEM micrographs were taken for three independent batches of LPAA of each lysine/valine ratio. Micrographs were then further analysed to determine LPAA particle size using Bruker ESPRIT 2.0 particle analysis software package.

**Scheme 1:**
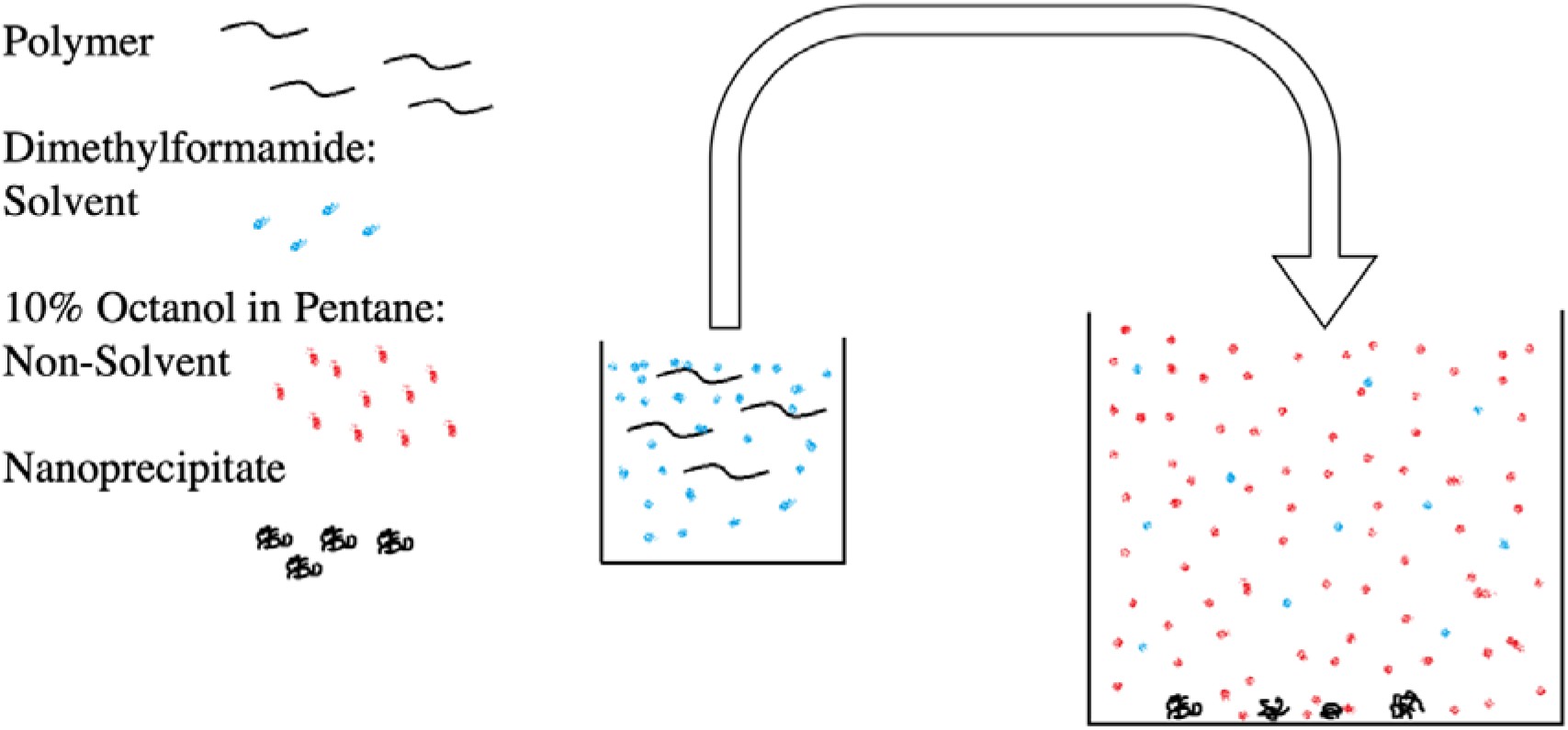
Nanoprecipitation process. LPAANPs were formed by first dissolving polymer in dimethylformamide (DMF) at 10 mg/mL and then adding the solution dropwise into a magnetically stirred solution of 10% octanol in pentane. As DMF diffuses into the octanol/pentane solution, LPAAs form nanoparticles (LPAANPs) which are collected via ultracentrifugation for further use.

### Nanoprecipitation

LPAAs were dissolved at 10 mg/mL in DMF (solvent) with the assistance of 5 minutes of sonication employing a bath sonicator (Fisher FS30). A mixture of 10% octanol and 90% pentane (nonsolvent) was prepared for use in the nanoprecipitation process. LPAA solution (500 μL in DMF) was added dropwise to 25 mL of octanol/pentane in a 50 mL ultracentrifuge tube under constant stirring. After 5 minutes of stirring, ultracentrifugation at 20,000 g for 20 minutes (Thermo Sorvall Lynx 6000) was used to sediment the LPAA nanoparticles (LPAANPs). After centrifugation, the solution was decanted and an additional 10 mL of pentane added to remove any extra DMF and octanol. Following another centrifugation cycle and decantation, the pentane rinse was repeated twice more for a total of three washes. LPAANPs were then dried under high vacuum for 1 hour and stored at room temperature in the dark until used. A depiction of the nanoprecipitation process is provided in **Scheme 1**.

### Fibroblast Toxicity of LPAAMs and LPAANPs

#### Fibroblasts Cell Culture and Treatment

3T3 fibroblasts (ATCC L929) were generously provided by Dr. Sheila Grant. These cells were cultured in Dulbecco’s Modified Eagle’s Medium (DMEM) supplemented with 10% fetal bovine serum (FBS) and penicillin-streptomycin using 75 cm^2^ cell culture flasks at 37°C in a humidified incubator at 5% atmospheric CO_2_ as per ATCC protocols. Media was changed every 3 days to remove waste and supply new nutrients. Fibroblasts were harvested near complete confluency for toxicity assays via trypsin delamination. Cells were counted and seeded in clear, sterile 96 well plates with 5,000 cells/well in 100 μL of media. Fibroblasts were further cultured for 48 hours prior to the addition of any material for toxicity testing. Stock solutions of each particle type (LPAAMs and LPAANPs) were prepared at 10000, 5000, 2000, 1000, 500, 100, and 0.0 μg/mL. Each treatment (11 μL) was added to incubating cells for final concentrations of 1000, 500, 200, 100, 50, 10 and 0.0 μg/mL in PBS and media and cultured for 24 hours. Following treatment, the media from each well was emptied with care taken to not disturb the adherent cells. Each well was rinsed twice with 100 μL PBS followed by the addition of 100 μL of 1% Triton X-100 in ddH_2_O. Cells were then subjected to three freeze-thaw cycles to lyse the cells and release their intracellular components. All cell processing was completed using proper sterile technique. Cell culture and treatment with each formulation was completed three times in triplicate.

#### DNA Assay

DNA content was measured to assess the number of 3T3 fibroblasts present after 24 hours of incubation with each experimental group using a Quant-iT™ PicoGreen dsDNA kit (Invitrogen). Following the aforementioned freeze-thaw cycles, 5 μL of each treatment group was added to 95 μL of pH 7.5 TE buffer (10 mM tris(hydroxymethyl)aminomethane and 1 mM ethylenediaminetetraacetic acid) in a black 96-well plate. Working PicoGreen solution (100 μL of 1x PicoGreen reagent in TE) was prepared fresh and added to each well. The plate was incubated in the dark at room temperature for 5 minutes followed by fluorescent quantification using an excitation/emission of 485/520 nm (BioTek Cytation5 spectrofluorometric plate reader). A DNA content / cell number standard curve was generated by lysis of a known concentration of cells (1,000,000 cells in 100 μL of 1% Triton-X100 in ddH_2_O) followed by serial dilution and DNA quantification using the same protocol as that for experimental samples. Each assay was completed three times in triplicate.

#### ATP Assay

ATP content was evaluated and compared to a standard curve to assess the metabolic health of 3T3 fibroblasts exposed to each experimental group. Each cell lysis sample (10 μL) was added to 90 μL of PBS to dilute the 1% Triton X-100 to 0.1%. Some of this solution (25 μL) was added to a 384-well white plate. 25 μL of Cellglo Buffer made from the CellTiter-Glo Luminescent Cell Viability Assay (Promega) was then added to each well. Samples were shaken for 2 minutes, the plate incubated at room temperature for 10 minutes, and then luminescence was measured with a BioTek Cytation5 specrofluorometric plate reader. Care was taken to prevent bubbles from forming during this process. An ATP standard curve was prepared by serial dilution of a 10 μM ATP standard from 1000 nM to 0.1 nM and luminescence quantified in parallel with experimental samples. Each assay was completed three times in triplicate.

### Peptide, Peptide Amphiphile, and Doxorubicin Entrapment

5(6)-Carboxyfluorescein (FAM)-labeled peptide (FAM-GRKKRRQRRRPPRPDRKLEVFEKEFLRMELGERC) and Fam-labeled peptide amphiphile (Palm_2_KK(FAM)-RPDRKLEVFEKEFLRMELGER) were synthesized on a Tetras peptide synthesizer (Advanced ChemTech) using a standard orthogonal fluorenylmethyloxycarbonyl (Fmoc) protection strategy with hexafluorophosphate benzotriazole tetramethyl uronium (HBTU)-mediated amino acid conjugations supplemented with hydroxybenzotriazole (HOBt) and N,N-diisopropylethylamine (DIEA). Fmoc deprotection was completed by treatment with 25% piperidine in DMF. For fluorescent labeling of the peptide, FAM was conjugated after HBTU activation to the N’ peptide amino group. For peptide amphiphile FAM labeling, orthogonal protection of a lysine side chain via 1-(4,4-dimethyl-2,6-dioxocyclohexylidene)ethyl (Dde) was used during Fmoc-Lysine(Fmoc)-OH and palmitic acid couplings. After palmitic acid coupling via HBTU activation, Dde was removed by 2% v/v hydrazine in DMF. FAM was then conjugated to the lysine side chain of the peptide amphiphile on resin using standard HBTU chemistry. Following on resin synthesis, cleavage was completed using TFA with scavengers (2.5% v/v of water, triisopropyl silane, thioanisole, and ethanedithiol supplemented with 2.5% w/w phenol) followed by diethyl ether precipitation. Peptide (P) and peptide amphiphile (PA) were purified to > 95% via mass spectrometry controlled high pressure liquid chromatography (LC/MS) using a Waters System Gold chromatograph equipped with an in-line mass spectrum analyzer (Thermo Orbitrap). Doxorubicin (Dox) was purchased from Selleck Chemicals, stored at −20°C in accordance with supplier recommendations, and used as received. Prior to nanoprecipitation, P, PA, and Dox were dissolved in DMF at 10 mg/mL. Each of these were individually mixed with a 10 mg/mL LPAA solution and immediately nanoprecipitated using the same protocol as previously stated. A weight ratio of 1:9 entrapped species : LPAA was used for each formulation. After nanoprecipitation and the first centrifugation cycle, a portion of the 10% octanol / 90% pentane supernatant solution was retained for assessment of co-precipitation yield. P-, PA-, and Dox-containing LPAANPs were processed post-nanoprecipitation identically to LPAANPs as previously described.

### Entrapment Efficiency and Release Characteristics Evaluation

Co-precipitation efficiency of P, PA, and Dox was evaluated by measuring the fluorescence of the nanoprecipitation supernatant. After the first centrifugation of the nanoprecipitate process, 250 μL of the supernatant was pipetted into a microcentrifuge tube, and the pentane allowed to evaporate off overnight in the dark. Supernatant solution (10 μL) was then added to 65 μL of DMSO. The fluorescence of the resultant solution was evaluated in a 96-well black plate at either excitation/emission of 495/520 nm (P and PA) or 500/600 nm (Dox). A BioTek Cytation5 spectrofluorometric plate reader was utilized in this and all following experiments for fluorescent quantification. The fluorescence of each sample was compared to standard curves of each material produced by the addition of P, PA, or Dox to a solution 10% octanol in pentane followed by overnight evaporation and serial dilution into DMSO, utilizing the same content as for the experimental samples. For Dox release efficiencies, 2.5 μL of nanoparticles of each doxorubicin-containing polymer was diluted from 1 mg/mL in PBS into 97.5 μL of DMSO. The fluorescence of 75 μL of this solution was measured and the value compared to standard curves of Dox produced in the same solvent (2.5% PBS in DMSO). Release kinetics in PBS were evaluated by suspending nanoparticles of each formulation with each entrapped species in PBS at 1 mg/mL. Aliquots from each solution were taken throughout incubation at 37°C. The released material from each aliquot was separated from the nanoparticles via separation using 50 kDa molecular weight cutoff filtration. The filtrate fluorescence was evaluated along with standards of each entrapped material in PBS. Aliquots were taken at 0, 3, 6, 12, 24, and 72 hours for P and PA and at 0, 24, and 72 hours for Dox. Entrapment efficiencies, Dox release efficiencies in DMSO, and release kinetics in PBS were each evaluated from three independent nanoprecipitation batches of each relevant formulation.

### Nanoparticle Surface Modification

#### Maleimide Surface Decoration

Dox-entrapped Poly(K_6_V_54_)C_16_ NPs (*i.e*. LPAANP_6/54_Dox) were surface modified to display cell-targeting aptamer. Maleimide-diethyleneglycol-tetrafluorophenol ester (Mal-DEG-TFP) was purchased from Quanta Biodesign and stored at −20 °C until use. Mal-DEG-TFP was dissolved in ddH_2_O at 10 mM with the aid of bath sonication for 10 minutes. LPAANP_6/54_ Dox (0.1 mg) was suspended in 100 mM sodium bicarbonate at a concentration of 1 mg/mL in a microcentrifuge tube. Mal-DEG-TFP (10 μL of the 10 mM stock solution in ddH_2_O) was then added, for a final concentration of 0.9 mM Mal-DEG-TFP. This resulted in a ratio of Mal-DEG-TFP : potential reactive amine sites of 1:1.17 assuming an average molecular weight of 7,000 Da per Poly(K_6_V_54_)C_16_. The reaction was allowed to proceed for 1 hour, followed by the addition of 500 μL of saturated ammonium sodium bicarbonate (ASB) (^~^ 1 M) on ice. After 10 minutes of incubation on ice, the solution was centrifuged at 20,000 g for 20 min to sediment the Mal-LPAANP_6/54_ Dox and for particle isolation from unreacted Mal-DEG-TFP and tetrafluorophenol. The supernatant was discarded and the pellet rinsed with two additional cycles of 500 μL saturated ASB addition, centrifugation, and decantation.

#### Surface decoration with antitail and aptamer

DNA antitail (A), C10.36-tail (NHL-specific aptamer)^14^, G24A-tail (point mutant specificity control for C10.36), and scApt-tail (or scDW4, a nontargeting DNA aptamer)^36^ were purchased from Integrated DNA Technologies with sequences listed in **Table S1**. Antitail was purchased with a hexyl-protected 5’ thiol. Each aptamer was purchased either un-modified or with a 5’ aminohexyl group, and all sequences were stored in TE buffer at −20 °C in the dark until used. Antitail purchased with a protected thiol (5’ Thiol Modifier C6 S-S) was deprotected by treatment with 20x excess tris(2-carboxyethyl)phosphine (TCEP) in 10x PBS at pH 7.4 followed by molecular-weight-cutoff filtration (MWCO 3KDa) and rinsing with ddH_2_O to form thiolated antitail (HS-antitail) as previously described^37^. With an estimated LPAA molecular weight of 7,000 Da, a molar ratio of 1:150 HS-antitail:LPAA was used. To 0.1 mg of Mal-LPAANP_6/54_ Dox containing 14.3 nmol LPAA, 0.095 nmol of HS-antitail was added. This ratio was used to maintain comparable aptamer valence to that of previously studied peptide amphiphile micelles displaying aptamer^37^. To assess whether additional functionalization density would improve LPAANP surface properties, HS-antitail was also added at 1:100 and 1:50 molar ratios to maleimide groups. For each case, HS-antitail was added to Mal-LPAANP_6/54_ Dox in PBS and allowed to react overnight. Lastly, aptamer-tail (C10.36, G24A, scApt, or Cy3 fluorophore-labeled derivatives of each aptamer), was annealed to A-Mal-LPAANP_6/54_ Dox. For fluorescent aptamer synthesis, Cy3-NHS ester was reacted with the amine end of each aptamer to produce Cy3-Apt as previously described^37^. In brief, a 20x excess of N-hydroxysuccinimide-Cy3 was reacted with each aptamer overnight followed by HPLC purification and then concentration and buffer exchange by molecular weight cutoff filtration. Each Cy3-aptamer was stored in pH 8.5 TE buffer in the dark at −20 °C until used. For aptamer decoration of 0.1 mg of A-Mal-LPAANP_6/54_ Dox, 0.29 nmol of aptamer (Apt) was added in PBS supplemented with 6.25 mM MgCl_2_ to aid in proper aptamer folding. A 3:1 Apt:A ratio was held and samples annealed by heating to 90 °C and cooling to room temperature over 45 minutes. Samples of each type (LPAANP_6/54_ Dox, Mal-LPAANP_6/54_ Dox, A-Mal-LPAANP_6/54_ Dox, Apt^~^A-Mal-LPAANP_6/54_ Dox, and Cy3-Apt^~^A/Mal-LPAANP_6/54_ Dox) were stored at 4°C in the dark at 500 μg/mL in PBS with 6.25 mM MgCl_2_ prior to further evaluation. A depiction of the steps of DoxNP surface modification is shown as **Scheme 2**.

#### TEM, Fourier-Transform Infrared Spectroscopy (FTIR), and Zeta Potential of Surface Modified Nanoparticles

Surface-modified nanoparticles were evaluated via TEM in a manner similar to that for non-modified LPAAMs and LPAANPs. However, due to the presence of DNA and PBS, these precipitates tended to be found within salt crystals. Care was taken to capture and evaluate representative micrographs of each particle type. FTIR spectra were captured in PBS at 500 μg/mL for each particle type utilizing a Nicolet 6700 FT-IR instrument (Thermo Scientific). Spectra were collected in the range of 3400 cm^−1^ to 400 cm^−1^. For zeta potential evaluation, surface-modified nanoparticles were diluted to 10 μg/mL in 2% PBS with ddH_2_O. Zeta potential measurements were taken using a Malvern Zetasizer Nano instrument. DTS1070 gold-plated zeta cuvettes were used with 700 μL of sample solution. Three measurements were taken per sample with standard manufacturer protocols followed. Surface-modified nanoparticles were evaluated from three independently produced batches.

**Scheme 2:**
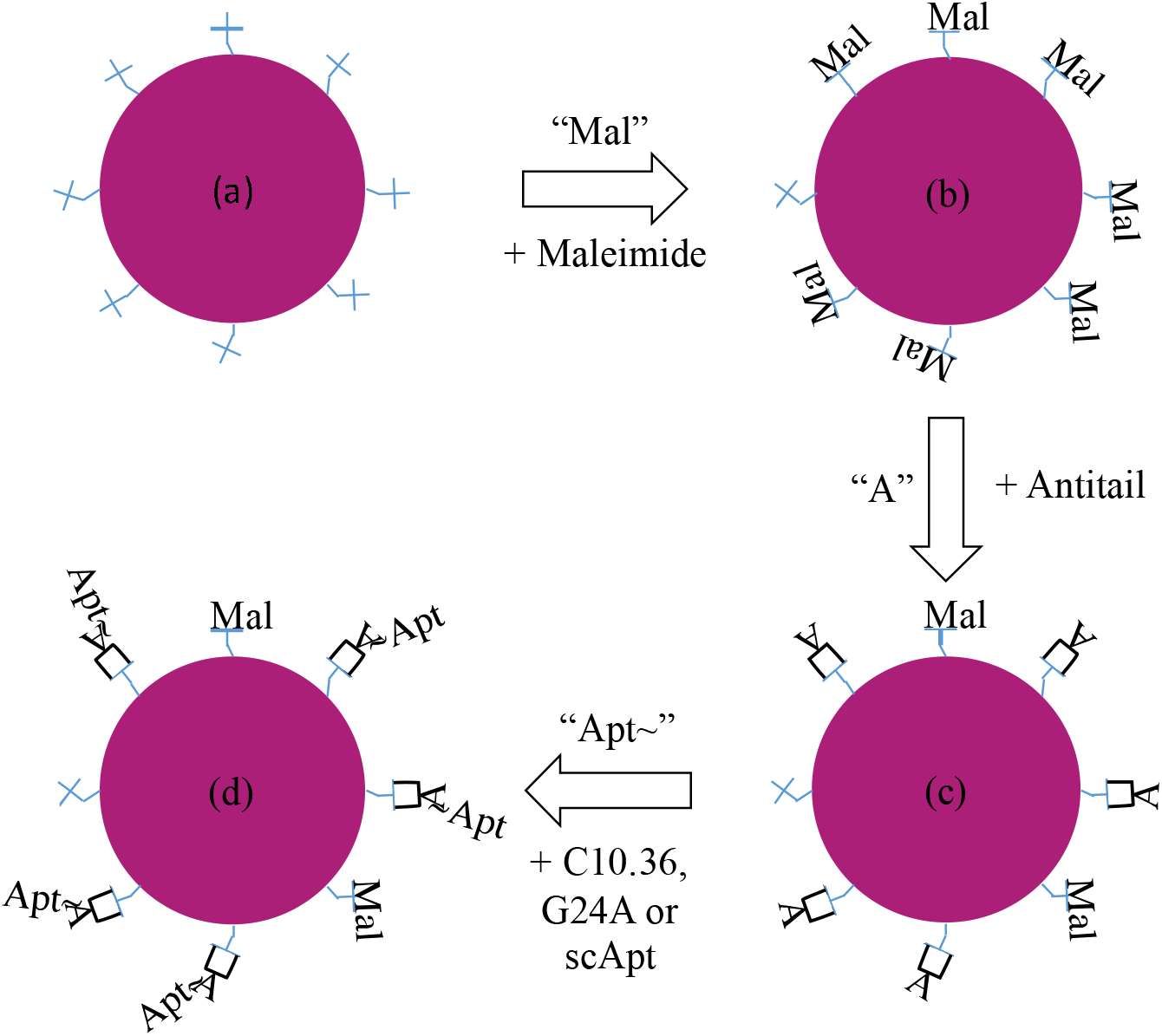
NP surface modification strategy. (a) Dox-entrapped Poly(K_6_V_54_)C_16_ NPs (*i.e*. LPAANP_6/54_ Dox) were prepared and suspended in phosphate buffered saline (PBS) with primary amines on the nanoparticle surface shown as blue crosses. (b) Mal-DEG-TFP ester was reacted with surface amines to yield Mal-LPAANP_6/54_ Dox, (c) thiolated antitail was reacted with surface maleimide to form A-Mal-LPAANP_6/54_ Dox, and (d) aptamer was annealed to surface antitail to generate Apt^~^A-Mal-LPAANP_6/54_ Dox.

### Cellular Delivery of Doxorubicin Via Surface Modified Nanoparticles

#### Aptamer-Mediated Nanoparticle Delivery

Surface-modified nanoparticles employing Cy3-aptamers were used for nanoprecipitate delivery experiments. To ensure that non-annealed aptamer was not present in the formulation, Cy3-aptamer was used at a sub-stoichiometric ratio (0.5:1) of Apt:A. Ramos cells (ATCC CRL-1596) were cultured in RPMI media supplemented with 10% FBS in a 37°C humidified incubator at 5% CO_2_. Prior to treatment, cells were centrifuged and counted. Suspensions of Ramos cells at 125,000 cells in 25 μL of 10% FBS in RPMI were prepared in a clear, round bottom 96-well plate. Salmon-sperm DNA (ssDNA, Sigma) was added (5 μL of 10 mg/mL) as a non-specific competitor for aptamer binding sites. Nanoparticles of each formulation at 250 μg/mL (20 μL) were added to the cell suspension for a total volume of 50 μL in each well. Each experimental group (Cy3-C10.36^~^A-Mal-LPAANP_6/54_ Dox, Cy3-G24A^~^A-Mal-LPAANP_6/54_ Dox, and Cy3-scApt^~^A-Mal-LPAANP_6/54_ Dox) was incubated with Ramos cells for 10 min followed by rinsing with 150 μL of 10% FBS in PBS and subsequent centrifugation at 1,400 RPM for 3 minutes (Thermo Sorvall Legend R+). The cells were then resuspended in 150 μL of 10% FBS in PBS and a second centrifuge cycle completed. The solution was then drawn off and 150 μL of 10% FBS in PBS was added and the cells were resuspended. Cells were then evaluated by flow cytometry utilizing a BD LSRFortessa X-20 instrument with at least 50,000 cells evaluated from each treatment group. Live lymphocytes were identified via forward- and side-scatter measurements. Cy3 fluorescence of live lymphocytes was quantified via calculating the geometric mean captured from the PE fluorophore channel. Sample plots of data acquisition and the gating strategy utilized can be found in **Figure S2**. This study was independently repeated three times. Aptamer-free samples such as LPAANP_6/54_ Dox, Mal-LPAANP_6/54_ Dox, and A-Mal-LPAANP_6/54_ Dox were not included because weak inherent fluorescence of entrapped Dox was found to be insufficiently bright for evaluation by flow cytometry (*data not shown*).

#### Doxorubicin-Dependent Ramos Cell Death

Modified nanoparticles (LPAANP_6/54_, C10.36^~^A-LPAANP_6/54_, LPAANP_6/54_ Dox, scApt^~^A-Mal-LPAANP_6/54_ Dox, G24A^~^A-Mal-LPAANP_6/54_ Dox, and C10.36^~^A-Mal-LPAANP_6/54_ Dox were prepared at 250 μg/mL in PBS. Free Dox was prepared at 25 μg/mL in PBS. All Dox-NPs formulations and free Dox were incubated with Ramos cell line (250,000 cells) for 10 minutes at 37°C in 100 μL that contained 50 μL of cell suspension in 10% FBS in RPMI, 40 μL of experimental sample, and 10 μL of ssDNA. This resulted in a final concentration of each nanoparticle formulation of 100 μg/mL and Dox concentration of 10 μg/mL. The concentration of Dox in each treatment with either free Dox or entrapped Dox were the same. Following the 10 minute incubation, 150 μL of 10% FBS in PBS was added to each well. Cells were then pelleted via centrifugation, supernatant decanted, and cell resuspended as previously described. After the second centrifugation cycle, cells were resuspended in RPMI supplemented with 10% FBS and cultured at 37°C for 24 hours. Cells were diluted with 100 μL of 10% FBS in PBS, centrifuged as before, and re-suspended in 100 μL of 10% FBS in PBS. 7-AAD stain (Invitrogen) was added to each sample (2.5 μL per well in the 96-well plate) and incubated for 10 minutes. Samples were immediately evaluated via flow cytometry to assess the percentage of dead cells compared to untreated cells. An example of the data acquisition and gating strategy is shown in **Figure S3**. Dead cells were identified via side-scatter versus forward-scatter measurements as well as from 7-AAD+ staining. 7-AAD+ cells were identified by plotting an empty channel (AlexaFluor 647, excitation/emission of 640/670 nm) by 7-AAD (excitation/emission of 488/710 nm). This study was independently repeated three times.

### Statistical Analysis

Comparisons between experimental groups were made using JMP software (SAS). A Tukey’s HSD test was completed for each pairwise analysis to establish whether there are any statistical differences between groups using an α value of 0.01. Experimental groups in each graph with statistically significant differences in means are noted with different letters.

## Results

### NCA polymerization produced well-defined LPAAs

Lysine(Cbz) and valine NCAs were synthesized and purified (**Figure S1**), after which cyclic monomer mixtures with lysine/valine NCA ratios of 30/30, 15/45, 6/54, and 3/57 were polymerized overnight to form LPAAs using hexadecylamine as an initiator (**Scheme S1**). NMR DOSY was employed to assess polymer diffusivity, which was scaled to molecular weight (**Table 1**). A representative NMR DOSY spectrum and the polystyrene standard curve used for diffusivity to molecular weight conversations are provided in **Figures S4** and **S5**, respectively. Additionally, a representative ^1^H NMR spectrum of the strategy used to determine lysine/valine ratios is provided in **Figure S6**. Measured molecular weights of protected LPAAs closely align with the expected values, as shown in **Table 1**. Additionally, the lysine/valine ratio of each LPAA follows a similar trend as the expected ratio. Each value shown is the average of three independently produced batches of LPAA for each formulation. All LPAAs were then deprotected using HBr in an acidic solution to remove the Cbz protecting groups. Deprotection was successful for each LPAA as confirmed via NMR, with a representative NMR provided in **Figure S7**.

**Table 1:** LPAA chemical characterization. Poly(K_a_V_b_)C_16_ with a variety of K:V ratios were synthesized with the expected linear structure at a 60:1 monomer:initiator ratio. Log diffusivity, lysine protected molecular weights, and actual lysine/valine ratios were determined by NMR DOSY and ^1^H NMR.

| LPAAs | Log Diffusivity (log<sub>10</sub>(m<sup>2</sup>/s)) | Measured Molecular Weight (Da) | Expected Molecular Weight (Da) | % Difference |
| --- | --- | --- | --- | --- |
| Poly(K<sub>30</sub>V<sub>30</sub>)C<sub>16</sub> | 9.51 ± 0.09 | 10300 ± 4000 | 11100 | - 7% |
| Poly(K<sub>15</sub>V<sub>45</sub>)C<sub>16</sub> | 9.45 ± 0.07 | 7500 ± 2200 | 8600 | -14% |
| Poly(K<sub>6</sub>V<sub>54</sub>)C<sub>16</sub> | 9.39 ± 0.12 | 5900 ± 3600 | 7200 | -27% |
| Poly(K<sub>3</sub>V<sub>57</sub>)C<sub>16</sub> | 9.44 ± 0.08 | 7200 ± 2400 | 6200 | +8% |

| LPAAs | Measured Lysine/Valine Ratio | Expected Lysine/Valine Ratio |
| --- | --- | --- |
| Poly(K<sub>30</sub>V<sub>30</sub>)C<sub>16</sub> | 0.95 ± 0.11 | 1 |
| Poly(K<sub>15</sub>V<sub>45</sub>)C<sub>16</sub> | 0.48 ± 0.09 | 0.33 |
| Poly(K<sub>6</sub>V<sub>54</sub>)C<sub>16</sub> | 0.20 ± 0.12 | 0.11 |
| Poly(K<sub>3</sub>V<sub>57</sub>)C<sub>16</sub> | 0.055 ± 0.022 | 0.053 |

### LPAAs spontaneousl form micelle (LPAAMs)

Poly(K_30_V_30_)C_16_, Poly(K_15_V_45_)C_16_, Poly(K_6_V_54_)C_16_, and Poly(K_3_V_57_)C_16_ each readily dispersed in PBS without the need for sonication or heating. Critical micelle concentration (CMC) experiments were completed to assess whether the LPAAs had self-assembled into micelles. All LPAAs possessed CMCs from 1.1 – 4.2 μg/mL (**Figure S8**) indicating LPAAM formation. To confirm the presence of these LPAAMs and assess their nature, each LPAAM was evaluated by TEM at 1 mg/mL in PBS, which is well above their respective CMCs. Representative micrographs are shown in **Figure 1**. Interestingly, while all LPAAs self-assembled into nanostructures, the more hydrophobic Poly(K_6_V_54_)C_16_ and Poly(K_3_V_57_)C_16_ yielded larger particles. Additionally, Poly(K_6_V_54_)C_16_ form liposomes, as seen by a double membrane via TEM, whereas Poly(K_3_V_57_)C_16_ forms large, solid core nanoparticles (**Figure S9**). Interestingly, scanning electron micrographs (SEM) failed to resolve discrete particles, perhaps due to inter-nanoparticle interactions and aggregation during sample drying (**Figure S10**).

**Figure 1:**
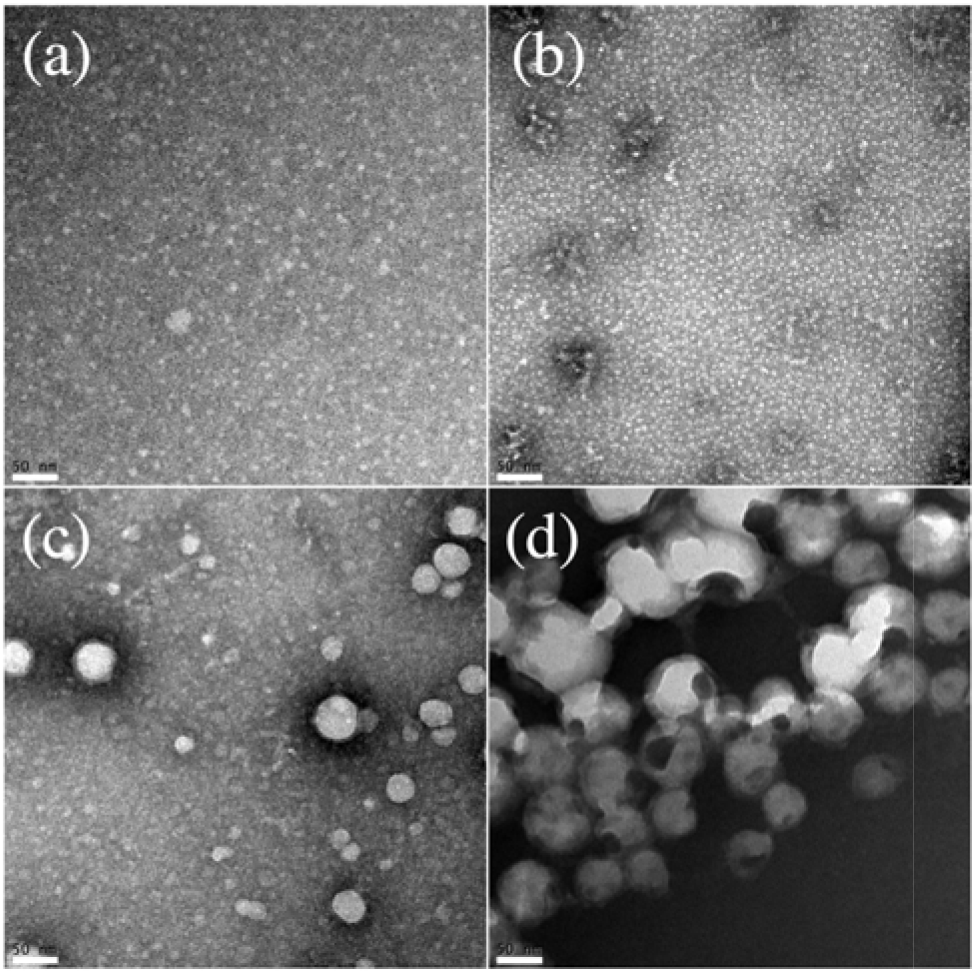
LPAA self-assembled particle structure. TEM micrographs showed all formulations - (a) Poly(K_30_V_30_)C_16_, (b) Poly(K_15_V_45_)C_16_, (c) Poly(K_6_V_54_)C_16_, and (d) Poly(K_3_V_57_)C_16_ - selfassemble in PBS into nanostructures (*i.e*., LPAAMs). Interestingly, the more hydrophilic LPAAs formed smaller nanostructures than the hydrophobic LPAAs (c) and (d). All scale bars are 50 nm.

### LPAAs can be readily nanoprecipitated

LPAAs were nanoprecipitated using a novel solvent/non-solvent system of DMF / 10% octanol in pentane. Each nanoparticle batch, termed LPAANP, was obtained in a good overall yield, with an average for all formulations of 78% ± 8%. An image size analysis program was utilized to assess the average diameter of each LPAAM and LPAANP batch whose protocol and example micrographs are provided (**Figure S11**). The compilation of this TEM-based data is shown in **Table 2**. LPAANPs were found to contain larger nanostructures than LPAAMs regardless of chemistry. The more hydrophilic LPAAs, Poly(K_30_V_30_)C_16_ and Poly(K_15_V_45_)C_16_, were found to yield nanostructures similar in small diameter regardless of fabrication method. The modestly hydrophobic Poly(K_6_V_54_)C_16_ formed small LPAAMs, but larger LPAANPs, whereas Poly(K_3_V_57_)C_16_, formed larger LPAAMs and LPAANPs.

**Table 2:** LPAAM and LPAANP size comparison. TEM micrographs of LPAA nanostructures were assessed by a TEM image analysis program to determine their size.

| LPAAs | Micelle Diameter (nm) | Nanoprecipitate Diameter (nm) |
| --- | --- | --- |
| Poly(K<sub>30</sub>V<sub>30</sub>)C<sub>16</sub> | 6.8 ± 3.6 | 12 ± 8 |
| Poly(K<sub>15</sub>V<sub>45</sub>)C<sub>16</sub> | 5.9 ± 3.0 | 17 ± 26 |
| Poly(K<sub>6</sub>V<sub>54</sub>)C<sub>16</sub> | 6.2 ± 5.4 | 79 ± 34 |
| Poly(K<sub>3</sub>V<sub>57</sub>)C<sub>16</sub> | 73 ± 21 | 190 ± 110 |

### LPAAMs and LPAANPs have promising toxicity profiles

Due to the interesting structural characteristics of LPAAMs and LPAANPs as potential drug delivery modalities, 3T3 murine fibroblasts were incubated with varying concentrations (10 – 1000 μg/mL) of each formulation for 24 hours. Any toxic effects of nanostructures on cells was determined via assessment of relative DNA and ATP content compared to untreated cells (**Figure 2**). DNA was converted to cell number utilizing an internally created standard to evaluate the impact nanostructure had on proliferation whereas ATP was used as a proxy for cell metabolism. As shown in **Figure 2(a)** and **2(c)**, both the number of cells and ATP concentration were reduced at 1000 μg/mL for Poly(K_30_V_30_)C_16_ Ms and at 500 and 1000 μg/mL for Poly(K_15_V_45_)C_16_ Ms. For LPAANPs, no toxic affect was observed for any chemistry up to 1000 μg/mL maximum concentration tested (**Figure 2(b)** and **2(d)**).

**Figure 2:**
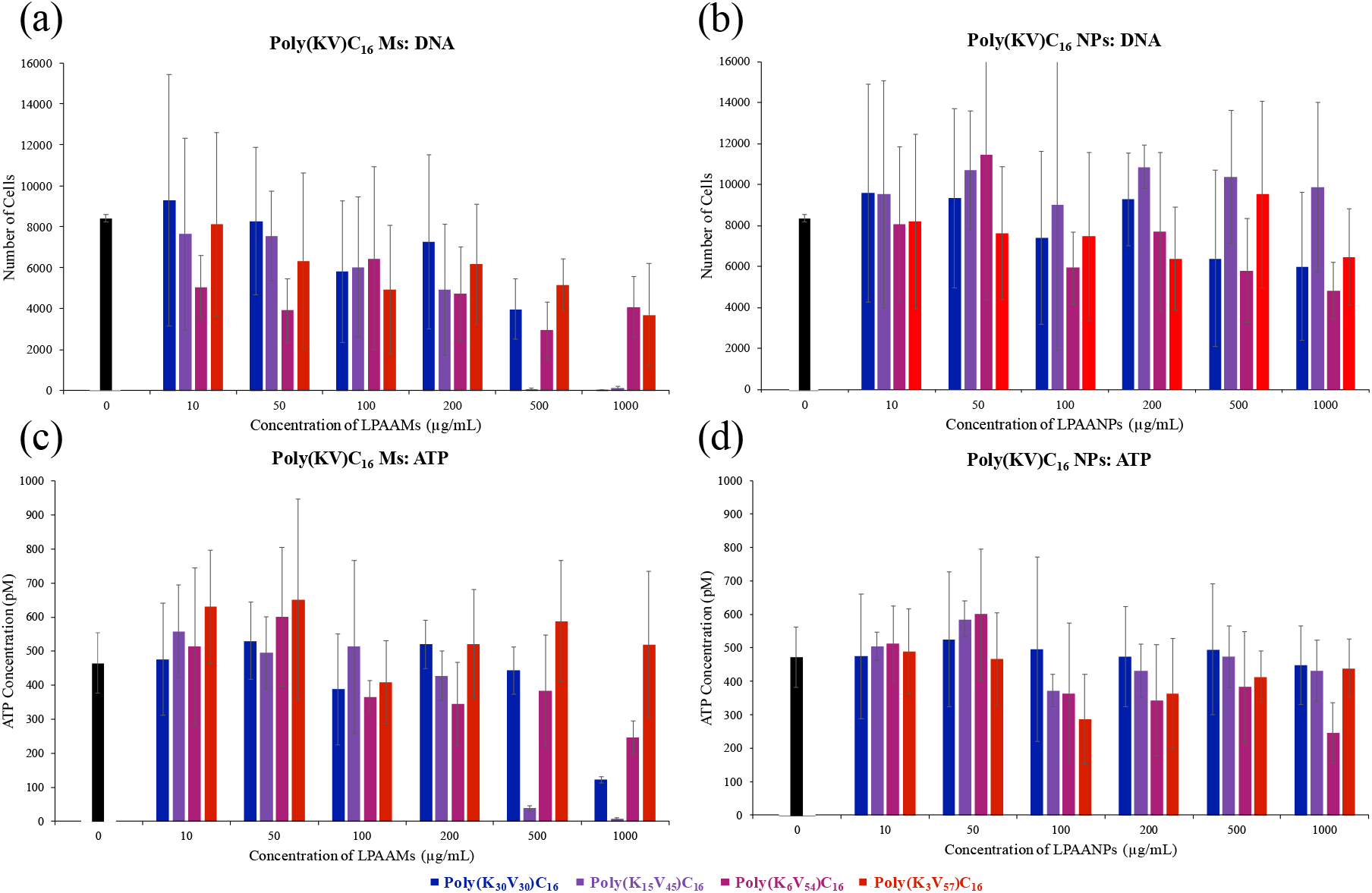
LPAAM and LPAANP cell toxicity. NIH 3T3 fibroblasts were treated with ((a) & (c)) LPAAMs or ((b) & (d)) LPAANPs for 24 hours to determine any formulation- and concentration-dependent toxicity. Cells were assayed for ((a) & (b)) DNA content and ((c) & (d)) ATP content for which no LPAAM formulation below 500 μg/mL and any LPAANP formulation tested negatively impacted cell health.

### LPAANPs efficiently entraps doxorubicin, peptide, and peptide amphiphile

The capacity for LPAANPs to be able to entrap doxorubicin (Dox), peptide (P), and peptide amphiphile (PA) was explored. LPAA and Dox, P, or PA were dissolved in DMF and precipitated in 10% octanol in pentane. After isolating the particles from solution, payload-nanoparticle association efficiency was assessed. Supernatant from the nanoprecipitation process were evaluated for either Dox fluorescence or FAM fluorescence as P and PA were fluorophore labeled. All drug payload association efficiencies were > 97% regardless of LPAA formulation chemistry and entrapped species (**Table 3**). Dox-, P-, and PA-loaded LPAANPs were then incubated in PBS to assess drug release kinetics. The immediate release, or burst release, of each payload from each LPAANP formulation is given in **Table 3**. Dox, P, and PA had burst releases of ^~^ 14% - 27%, P of ^~^ 47% - 72%, and PA of ^~^ 0.1% −10%, respectively. Over 72 hours of incubation in PBS, no further release beyond this burst was observed for Dox and P (**Figure S12**). Interestingly, PA did slowly reach ^~^ 40% release over three days for two formulations (Poly(K_30_V _30_)C_16_ NPs and Poly(K_3_V_57_)C_16_ NPs), whereas no extended payload release kinetics were found for Poly(K_15_V_45_)C_16_ NPs and Poly(K_6_V_54_)C_16_ NPs. LPAANP Dox were chosen as a promising candidate for drug delivery, so LPAANP Dox were then diluted into DMSO to dissolve the NPs and establish Dox release efficiencies from these particles, as given in **Table S2**. Dox release efficiency differed for each LPAANP and was found to be 92 ± 12% for Dox-loaded Poly(K_6_V_54_)C_16_ NPs (LPAANP_5/64_ Dox). This formulation was then surface modified with an NHL-targeting aptamer for further development.

**Table 3:** LPAANP drug association efficiency and burst release. LPAAs were co-precipitated with doxorubicin, peptide, and peptide amphiphile to evaluate entrapment efficiency and burst-release profiles for a wide range of potential drug classes.

| LPAANP | Associated Species | Association Efficiency | % Burst Release of Entrapped Species |
| --- | --- | --- | --- |
| Poly(K<sub>30</sub>V<sub>30</sub>)C<sub>16</sub> NP | Doxorubicin | 97.8 ± 0.2% | 15 ± 9% |
|  | Peptide | 100.1 ± 0.1% | 47 ± 13% |
|  | Peptide amphiphile | 99.9 ± 0.3% | 10 ± 1% |
| Poly(K<sub>15</sub>V<sub>45</sub>)C<sub>16</sub> NP | Doxorubicin | 97.6 ± 0.2% | 24 ± 14% |
|  | Peptide | 100.0 ± 0.1% | 63 ± 21% |
|  | Peptide amphiphile | 100.1 ± 0.2% | 1.2 ± 0.7% |
| Poly(K<sub>6</sub>V<sub>54</sub>)C<sub>16</sub> NP | Doxorubicin | 98.3 ± 0.3% | 14 ± 8% |
|  | Peptide | 100.0 ± 0.0% | 49 ± 22% |
|  | Peptide amphiphile | 100.0 ± 0.1% | 0.1 ± 0.4% |
| Poly(K<sub>3</sub>V<sub>57</sub>)C<sub>16</sub> NP | Doxorubicin | 99.0 ± 0.1% | 27 ± 7% |
|  | Peptide | 99.9 ± 0.1% | 72 ± 17% |
|  | Peptide amphiphile | 99.9 ± 0.3% | 5 ± 3% |

### Dox-loaded LPAANPs can be readily surface-modified to display aptamer

For use in the delivery of its doxorubicin cargo, LPAANP_6/54_ Dox nanoparticles were modified to display a cell-specific aptamer. Maleimide-DEG-TFP ester was reacted with surface amines on LPAANP_6/54_ Dox to form Mal-LPAANP_6/54_ Dox. The presence of the maleimide group and change in surface amines from primary to secondary was indicated via FTIR (**Figure S13**). Antitail, the DNA oligomer with a thiol function group, was then added to react with maleimide and form A-Mal-LPAANP_6/54_ Dox with a variety of antitail:lysine ratios. Finally, aptamer, such as cell-targeting C10.36 that bears the tail sequence, was annealed to the surface to form Apt^~^A-Mal-LPAANP_6/54_ Dox. The change in zeta potential after each modification is shown in **Figure 3**. Additionally, TEM was utilized to evaluate any gross structural changes from surface modification. No differences were seen for each surface modification compared to unmodified LPAANP_6/54_ Dox (**Figure S14**).

**Figure 3:**
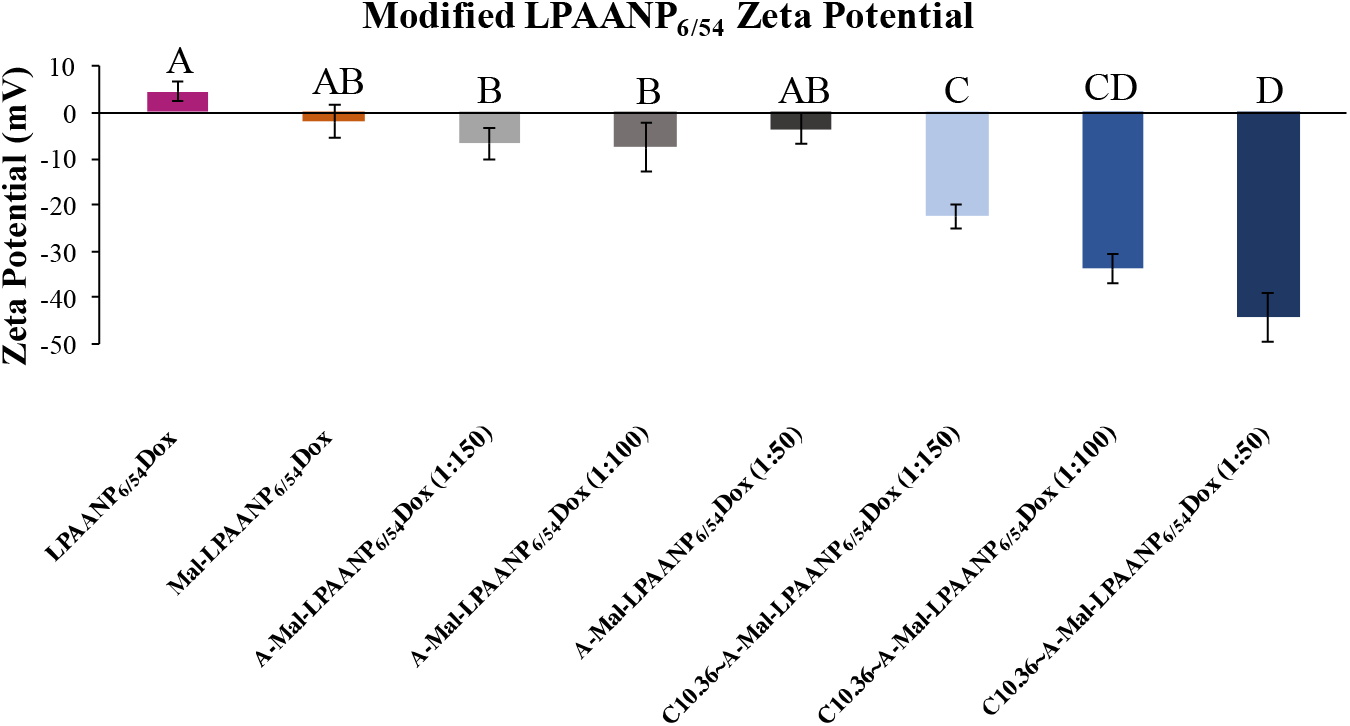
LPAANP surface modification impact on zeta potential. LPAANP_6/54_ Dox was sequentially surface modified to display cancer cell targeting aptamer (*i.e*. C10.36). The zeta potential was found to decrease after the conjugation of maleimide and antitail as well as the annealing of aptamer. The concentration-dependent manner of this behavior indicates desirable variance in the quantity of surface-displayed aptamer. Groups with different letters have statistically significant differences in means (Tukey HSD, ⍰ = 0.01).

### Aptamer-modified LPAANPs achieve enhanced cancer cell death

To test the *in vitro* functionality of the aptamer after annealing, Ramos cells were treated with Cy3-Apt^~^A-Mal-LPAANP_6/54_ Dox. After a 10 minute incubation with each treatment, cells were rinsed and evaluated by flow cytometry. Relative Cy3 fluorescence was used as a measure for nanoparticle delivery to the cell. The mean fluorescent intensities from each sample compared to untreated cells are shown in **Figure 4**. ScApt and G24A were found to have a minor increase in fluorescence over background. However, C10.36-labeled NPs had a ^~^10-fold increase over background fluorescence, a statistically significant, aptamer-specific level of association to Ramos cells. This level of specificity is similar to that seen previously by *Opazo, et al.^14, 38^*. To determine whether this aptamer-dependent association would drive a biologically relevant delivery of LPAANP and subsequent release of Dox, Ramos cells were incubated for 10 minutes with Apt^~^A-Mal-LPAANP_6/54_ Dox and other formulations. The cells were rinsed and cultured for 24 hours at 37°C. Via flow cytometry, the toxicity of each experimental group was measured by 7-AAD death staining. As shown in **Figure 5**, nanoparticles without entrapped doxorubicin had a similar level of death as untreated cells. Additionally, aptamer C10.36 did not influence the background toxicity of doxorubicin-free LPAANP (C10.36^~^A-LPAANP_6/54_). C10.36^~^A-Mal-LPAANP_6/54_ Dox were more toxic to Ramos cells compared to all groups other than Dox, including equimolar free Dox.

**Figure 4:**
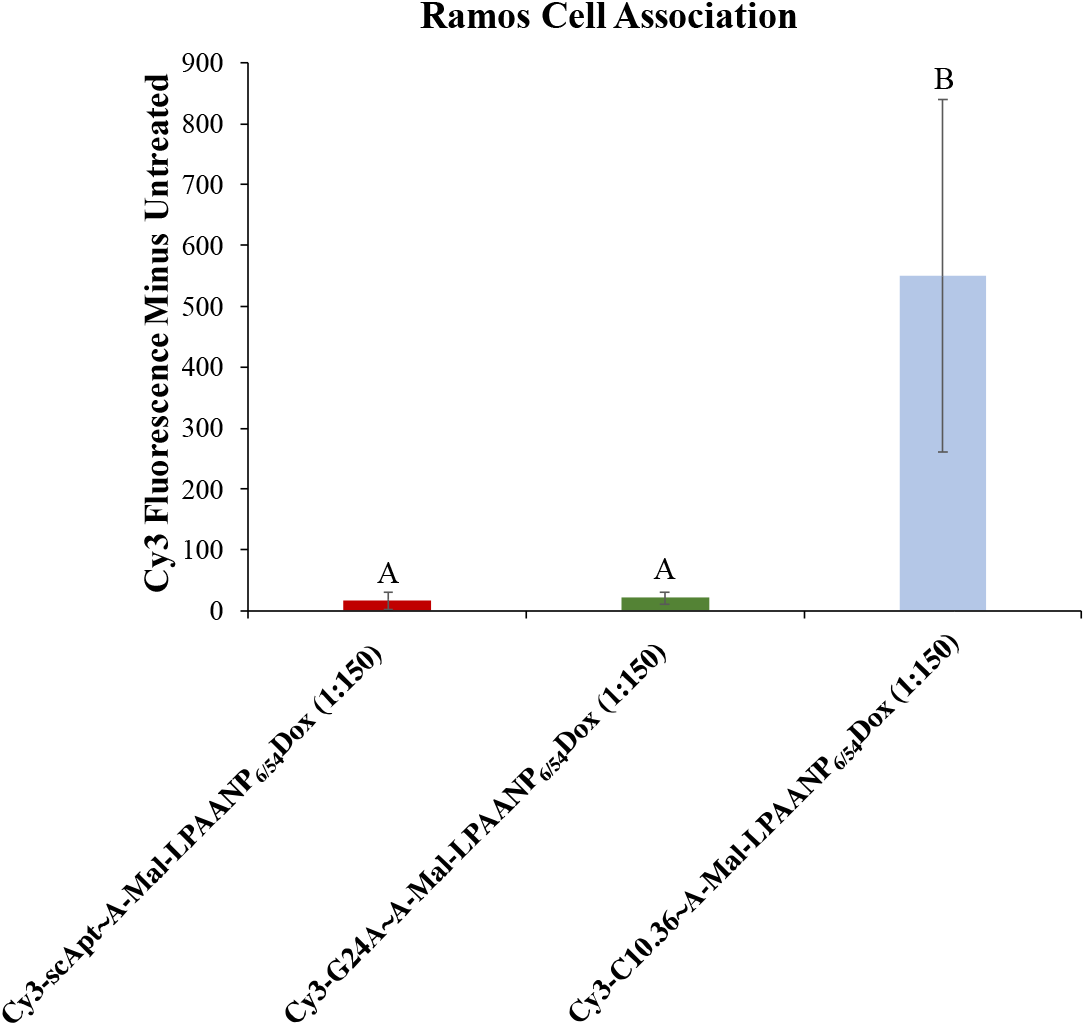
Surface-modified LPAANP aptamer specificity. Human lymphoma Ramos cells were incubated with doxorubicin-loaded, Cy3-labeled aptamer displaying nanoparticles (Apt^~^A-Mal-LPAANP_6/54_ Dox) for 10 minutes in the presence of 1 mg/mL ssDNA and 10% FBS in PBS. Following rinsing with 10% FBS in PBS to remove non-associated nanoparticles, cells were directly evaluated by flow cytometry. Cy3 fluorescence over untreated cells is shown which clearly shows aptamer-specific (*i.e*., C10.36-dependent) cellular association. Groups with different letters have statistically significant differences in means (Tukey HSD, ⍰ = 0.01).

**Figure 5:**
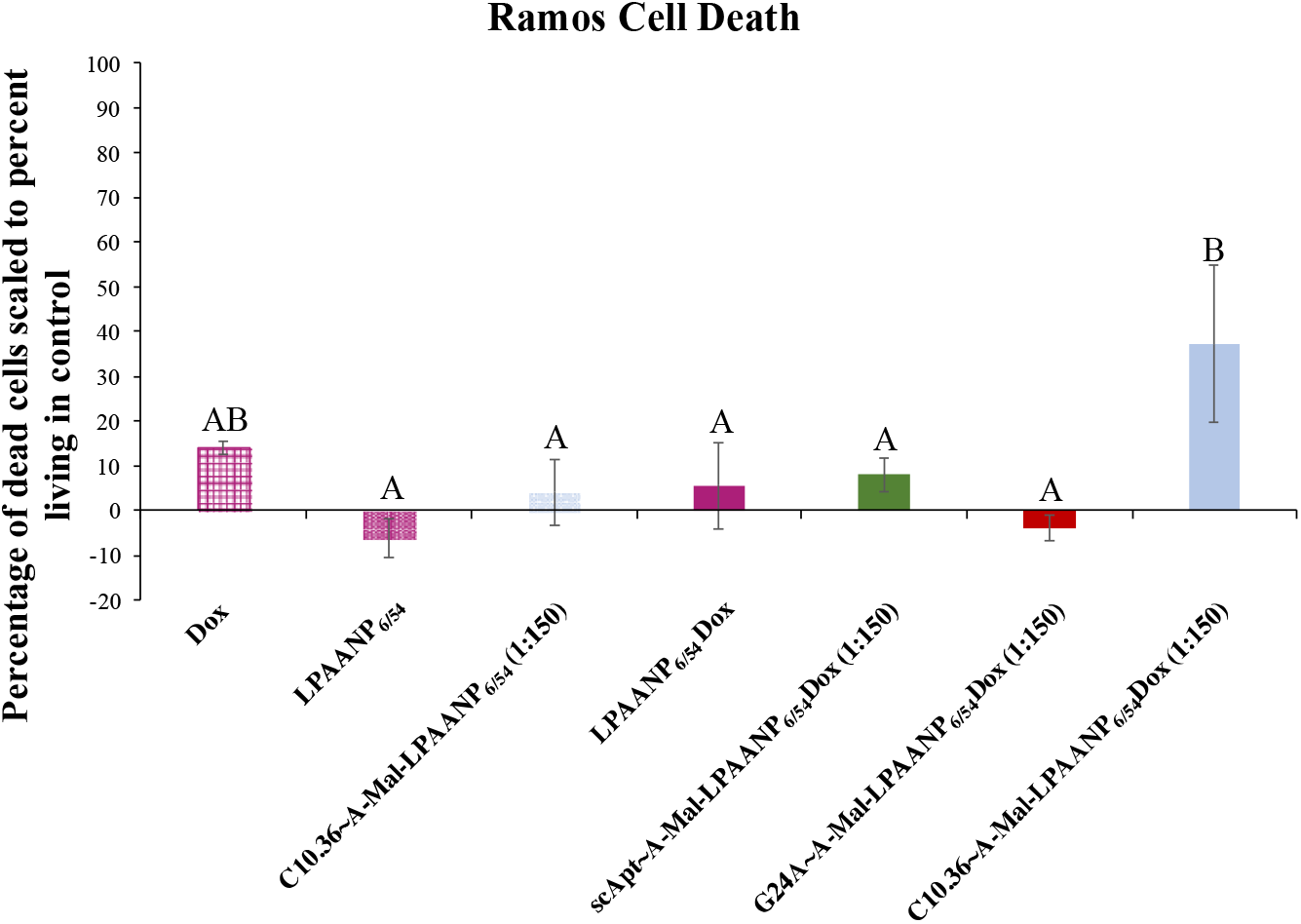
LPAANP-mediated Ramos cell toxicity. All formulations were incubated for 10 minutes, rinsed, and cultured for 24 hours. Cell death was then investigated using flow cytometry employing 7-AAD+ staining with data indexed against the population of untreated living Ramos cells. All Dox-containing formulations were utilized at a final Dox concentration of 10 μg/mL (17.2 μM). Groups with different letters have statistically significant differences in means (Tukey HSD, ⍰ = 0.01).

## Discussion

Poly(amino acid)s are a class of polymers unique in their facile synthesis, diverse amino acid composition, and straightforward incorporation of functional groups while remaining highly biocompatible and capable of drug delivery^22, 23, 39, 40^. In this research, hexadecylamine-initiated poly(amino acid)s that contain lysine and valine in various ratios (lysine/valine of 30/30, 15/45, 6/54, and 3/57) were produced, inspired by PAAs from *Lam et al.^41^*.

Each LPAA readily dispersed in PBS and spontaneously formed nanostructures, despite the hydrophobicity of these polymers with predominantly valine content. Based on the previously established properties of peptide amphiphiles, the hexadecyl group was expected to be a sufficiently long lipid chain to drive micellization^42–44^. However, beyond micellization, larger spontaneously forming nanostructures were observed (**Figure 1** and **Table 2**). These LPAA nanostructures may be formed to shield valine hydrophobic domains of each LPAA by aggregating and displaying primary amines from the lysine sidechains on their surface, similarly to other, more hydrophilic LPAAs previously reported^45, 46^. An increase in nanostructure size was observed from the more hydrophilic (Poly(K_30_V_30_)C_16_ and Poly(K_15_V_45_)C_16_) to the more hydrophobic (Poly(K_6_V_54_)C_16_ and Poly(K_3_V_57_)C_16_) LPAAs. This may be caused by the fewer lysine groups in Poly(K_6_V_54_)C_16_ and Poly(K_3_V_57_)C_16_ LPAAMs requiring a larger number of polymer chains to aggregate in order to provide a sufficient number of amine groups to stabilize the water-exposed surface of each nanoparticle.

To expand the utility of LPAAs beyond LPAAMs and to entrap drugs within LPAANPs, we developed a process for preparing LPAANPs by dissolving LPAA and P, PA, or Dox in DMF and precipitating via a dump-and-stir method into octanol/pentane, shown in **Scheme 1**. This method was found to be efficient for co-precipitating each LPAA and drug studied (**Table 3**). By comparison, several nanoprecipitation methods found in the literature may be applied to biomaterial systems like LPAAs to form NPs. The most similar example is dissolving drug and polymer in DMSO and using dialysis to controllably remove DMSO^31, 47, 48^. Additionally, aqueous methods have been published including various additives such as metals for chelation ^26, 49–51^. Advantages of this DMF into octanol/pentane process are the high precipitation efficiencies (>97%), biocompatibility of each solvent, ease of solvent removal, and short time required to produce nanoparticle batches. Furthermore, one concern in the development of a new LPAA system for drug delivery is processing-related and vehicular toxicities. As shown in **Figure 2**, two LPAAMs have a level of toxicity to fibroblasts at a dose of 500 μg/mL and no toxic effect below that concentration. Interestingly, LPAANPs had no toxic effect on fibroblasts at any concentration measured, which may be a result of altered nanostructure after nanoprecipitation.

Importantly, LPAANPs have shown tunability to appropriate size-ranges for drug delivery while optimizing loading efficiency of P, PA, and Dox (**Tables 2 and 3)^52^**. As a class of nanoprecipitate, one interesting aspect of these hydrophobic LPAANPs is the expected hydrolytic stability of the structure during blood stream circulation over commonly utilized, hydrolytically-sensitive polymer nanoparticles^53, 54^. With peptidase degradation of LPAANPs as the primary expected degradation route in biological systems, entrapped drug release may be delayed until cellular digestion via peptidases. This delay is advantageous by potentially minimizing off-target toxicity by reducing spontaneous drug release. Due to the small Dox burst release (14 ± 8%) and moderate size (79 ± 34 nm) of LPAANP_6/54_ Dox, this NP was further modified for the cellular delivery of entrapped doxorubicin. The surface amine functional groups from lysine were readily used for the attachment of targeting aptamer after maleimide and antitail functionalization (**Figures 3 and S13**). The resulting Apt^~^A-Mal-LPAANP_6/54_ Dox had an appropriate size and anionic surface charge for drug delivery in the blood stream (**Figures 3 and S14**)^52, 55^.

Aptamer C10.36 utilized in this Apt^~^A-Mal-LPAANP_6/54_ Dox formulation had previously been designed to selectively interact with lymphoma and leukemia cells, including NHL, and was previously used in the delivery of both RNA and peptide amphiphile micelles^14, 15, 37^. Excitingly, this high level of aptamer specificity to Ramos cells from a 10-minute incubation was retained after functionalization to the nanoparticle surface *in vitro* (**Figure 4**). Furthermore, this aptamer specificity was found to drive the delivery of Dox entrapped within LPAANP_6/54_ Dox to Ramos cells from a 10 minute incubation followed by 24 hour culture. A therapeutic level of Dox was released by C10.36^~^A-Mal-LPAANP_6/54_ Dox, outperforming non-specific aptamer formulations, as shown in **Figure 5**. Additionally, C10.36^~^A-Mal-LPAANP_6/54_ Dox were more toxic than all groups except for free Dox to Ramos cells. The aptamer-specific association and controlled toxic effect demonstrate the utility of LPAANPs as a class of polymer nanoprecipitates capable of delivery of a therapeutic molecule. This novel aptamer-specific nanoparticle delivery system for Dox is an original use of aptamer technology and exciting advancement in targeted therapeutics.

## Conclusion

The goal of this research was to explore the physical properties and drug delivery potential of a recently developed class of polymers, lipidated poly(amino acid)s (LPAAs). The controllability and scalability of the described NCA-derived LPAAs has attracted the attention of the research community, resulting in a variety of synthetic approaches for their generation as well as applicability in a variety of fields. In our research, we synthesized a series of poly(amino acid)s composed of lysine and valine. A nanoprecipitation method employing a novel DMF in octanol/pentane solvent system was used to fabricate nanostructures that could entrap a variety of small molecule, peptide, and peptide amphiphile therapeutic payloads. These drug delivery vehicles (LPAA nanoparticles, LPAANPs) were further altered through surface modification with an aptamer (*i.e*., C10.36) to facilitate their potential for cell-specific targeting. Excitingly, a chemotherapeutic-loaded, aptamer-displaying nanoparticle formulation (*i.e*., C10.36^~^A-Mal-LPAANP_6/54_ Dox) was found to associate with and induce the death of a human lymphoma cell line *in vitro*. Taken together, these data support the considerable potential of LPAANPs to function as a novel drug delivery vehicle platform for which encapsulated payload and cell targeting moiety can be modified for a variety of intended biomedical applications. Future work related to this project will focus on generating LPAAs incorporating a variety of other NCA amino acids (*e.g*., histidine and glutamic acid) and exploring the capacity for LPAANPs to selectively deliver anti-cancer peptide and peptide amphiphile payloads.

## Supporting information

Complete Supplementary File

## Acknowledgements

This work was supported by funds to BDU from the University of Missouri System Research Board and BioNexus KC as well as startup funds from the University of Missouri, to DHB from NIH award R21 AI121938, and to MAD from the University of Missouri System Research Board. We would like to thank the staff at the MU Electron Microscopy Core Facility, especially David Stalla, for their assistance in SEM imaging as well as image analysis. Additionally, we thank Dr. Sheila Grant for the use of her research group’s FTIR spectroscope and donating the NIH 3T3 fibroblasts.

