## Supplementary material for "Lipidated poly(amino acid) nanostructures as versatile therapeutic delivery vehicles": Complete Supplementary File

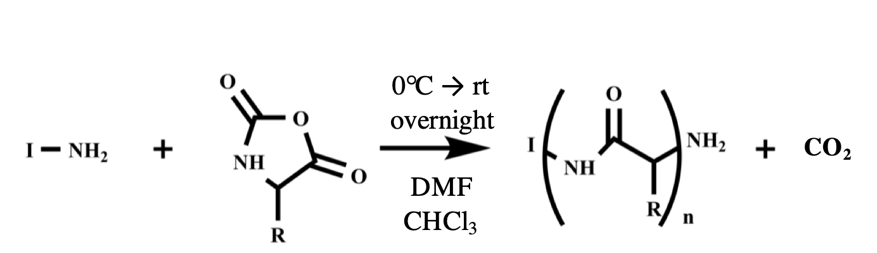

**Scheme S1:** N-carboxyanhydride (NCA) polymerization reaction. Any primary amine can act as an initiator (*I*) through which a NCA monomer ring is destabilized and attached followed by decarboxylation yielding a chain end primary amine group. This reaction propagates itself one NCA at a time until all monomer is incorporated into polymer strands. The reaction can be controllably carried out using a solvent mixture of dimethylformamide (DMF) and chloroform (CHCl_3_) warming from 0 ºC to room temperature (rt) overnight. For this research, hexadecylamine was used as an initiator.

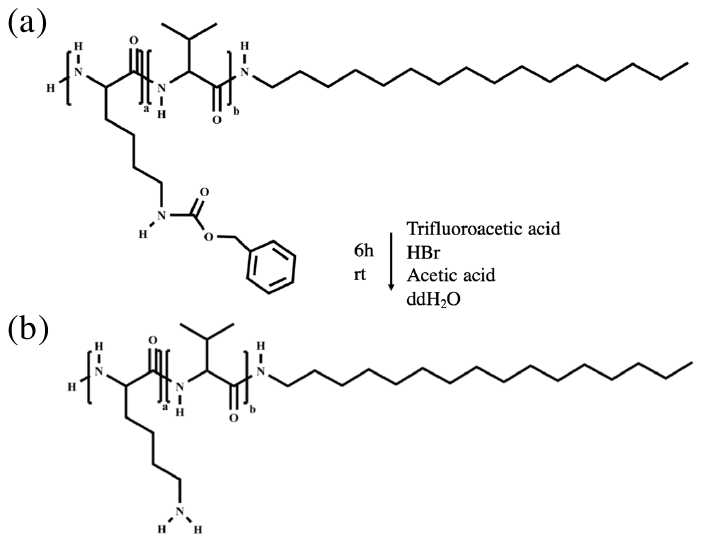

**Scheme S2:** Lipidated poly(amino acid) lysine side group deprotection. The carboxybenzyl (Cbz) group on lysine was deprotected to form poly(K(NH_2_)_a_V_b_)C_16_ by hydrobromic acid (HBr) treatment in a mixture of trifluoroacetic acid, acetic acid, and distilled, deionized water (ddH_2_O). This reaction was allowed to proceed for 6 hours at room temperature, followed by precipitation, neutralization, and lyophilization.

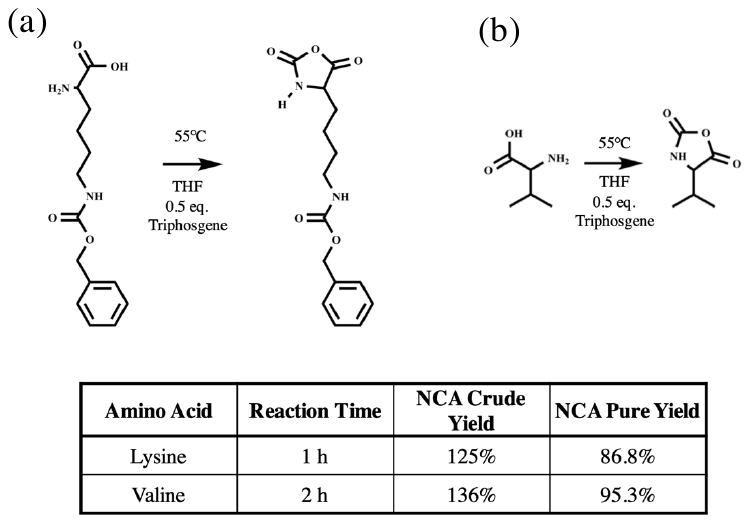

**Figure S1:** NCA synthesis and purification. (a) Lysine(Cbz) and (b) valine NCAs were synthesized as shown using the reaction time stated in the table. Crude and pure yields were determined by weight before and after purification by silica gel column chromatography. Lysine(Cbz) and Valine NCAs were successfully produced and confirmed by ^1^H NMR.

**Figure S2:** Cell association flow cytometry data analysis. (a) Dead cell and lymphocyte populations were identified by side scatter / forward scatter population evaluation. (b) PE histograms show the fluorescent intensity of cell associated Cy3-aptamer-labeled nanoprecipitates for which MFIs are reported in the table along with corresponding color coding and sample names.

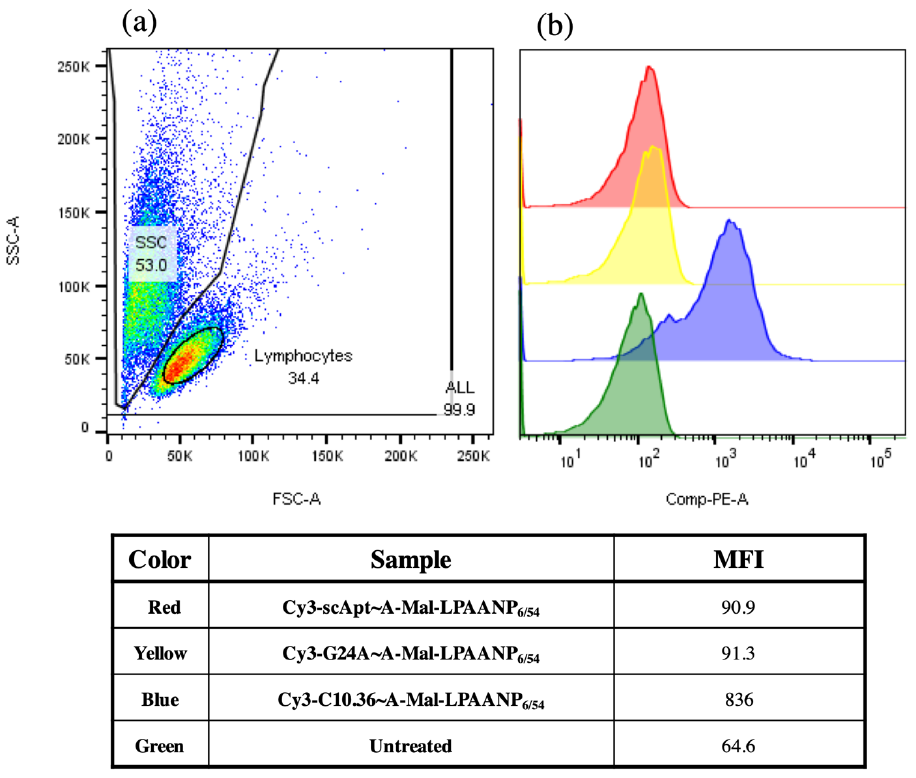

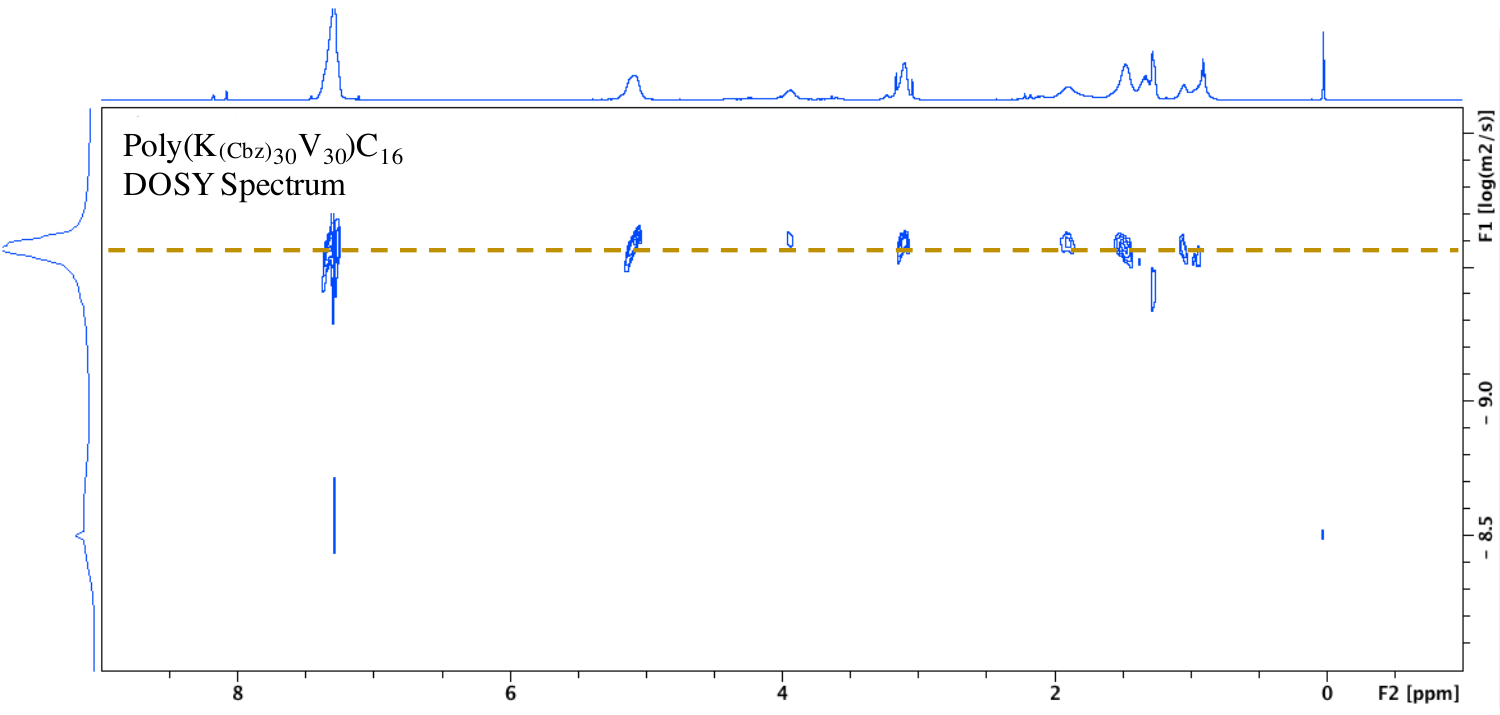

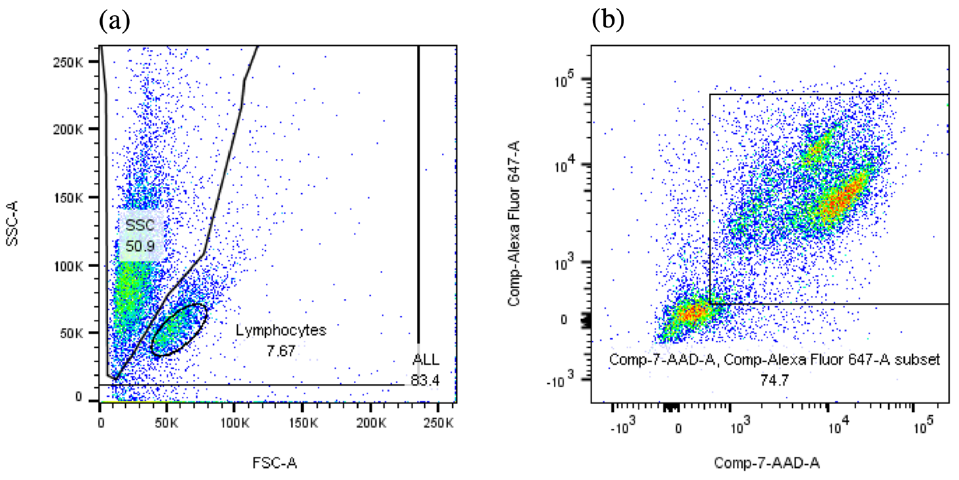

**Figure S3:** Cell death flow cytometry data analysis. (a) Dead cell were identified by side scatter / forward scatter population evaluation and (b) confirmed via 7-AAD staining. The results from one representative flow cytometry analysis are shown.

**Figure S4:** LPAA ^1^H NMR DOSY. A representative DOSY spectrum is shown for poly(K(Cbz)_30_V_30_)C_16_. The y-axis displays signal strength as the diffusivity (log(m^2^/s)) whereas the x-axis shows the NMR spectral shift. By selecting peaks containing the expected functional groups of the protected LPAA on the x-axis, the important diffusion coefficients can be readily identified using the y-axis.

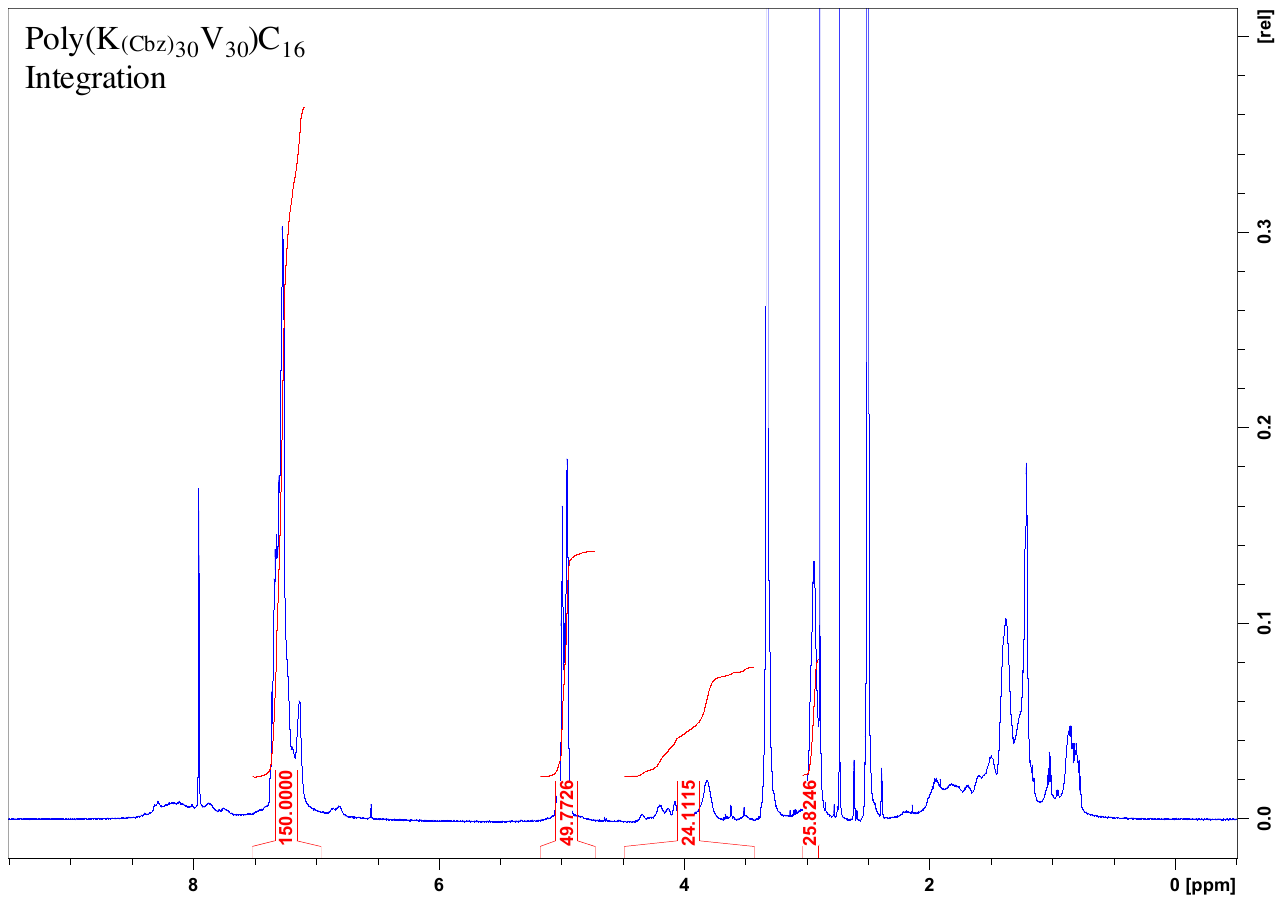

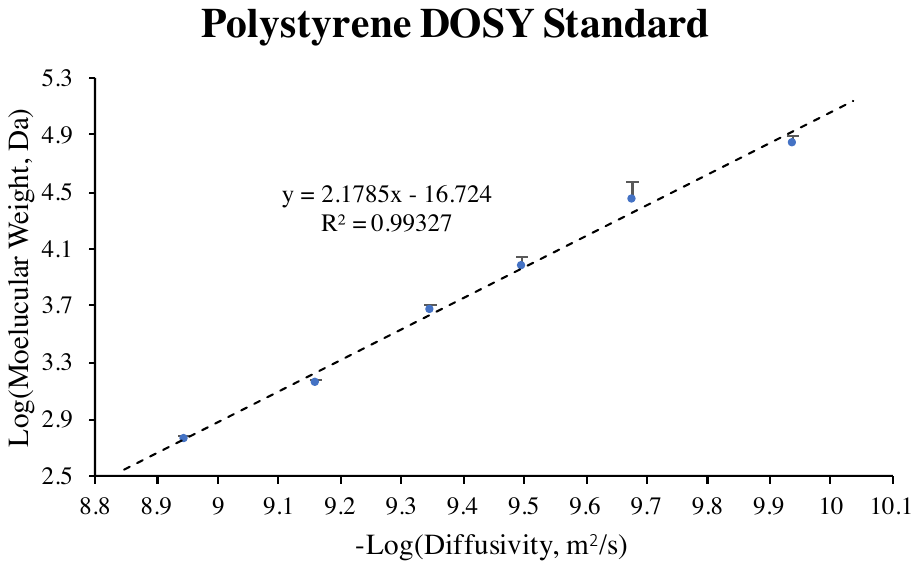

**Figure S5:** ^1^H NMR DOSY polystyrene standard curve. Uniform polystyrene standards were used to establish the relationship between diffusion coefficient and molecular weight in the solvent system utilized (*i.e.* 2.5% d-TFA in CDCl_3_). Inverse log of diffusivity and log of molar weight were found to be linearly correlated. Each data point shown was created from three independently prepared samples.

**Figure S6:** LPAA ^1^H NMR integration. The lysine/valine ratio was measured by integrating NMR spectra at 10 mg/mL sample in deuterated dimethyl sulfoxide (DMSO-d6). The Cbz peak (*i.e.* 4–, 5–, 6–, 7–, and 8–C hydrogens) at 7 – 7.5 ppm was set to an integration value of 150 to standardize plots against one another. The peaks from 3.5 – 4.5 ppm and at 3 ppm were attributed to the ⍺-carbon backbone hydrogens of lysine and valine, respectively, whose integration was used to estimate the calculated lysine-to-valine ratio.

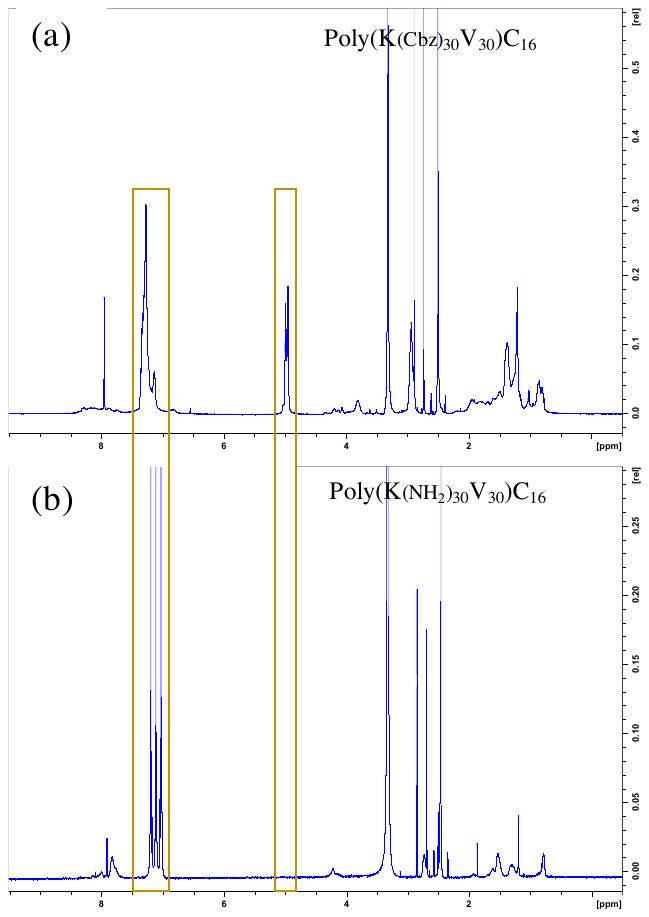

**Figure S7**: LPAA lysine(Cbz) deprotection confirmation. Sample ^1^H NMR spectra generated from LPAAs in DMSO-d6 (a) before and (b) after Cbz deprotection are shown. Characteristic peaks show the expected chemistry of (a) Poly(K(Cbz)_30_V_30_)C_16_ and (b) Poly(K(NH_2_)_30_V_30_)C_16_ as anticipated. Specifically, the disappearance of the singular 2-C hydrogen peak of Cbz at 5 ppm and changes in the 4–, 5–, 6–, 7– and 8–C hydrogen peaks at 7 – 7.5 ppm from that of Cbz (a single peak) to that of toluene (three discrete peaks) indicating successful deprotection. For some LPAAs, the post-deprotection toluene peak was not observed and instead no peaks were found at 7 – 7.5 ppm. In constrast, the disappearance of the 5 ppm peak was seen in all cases.

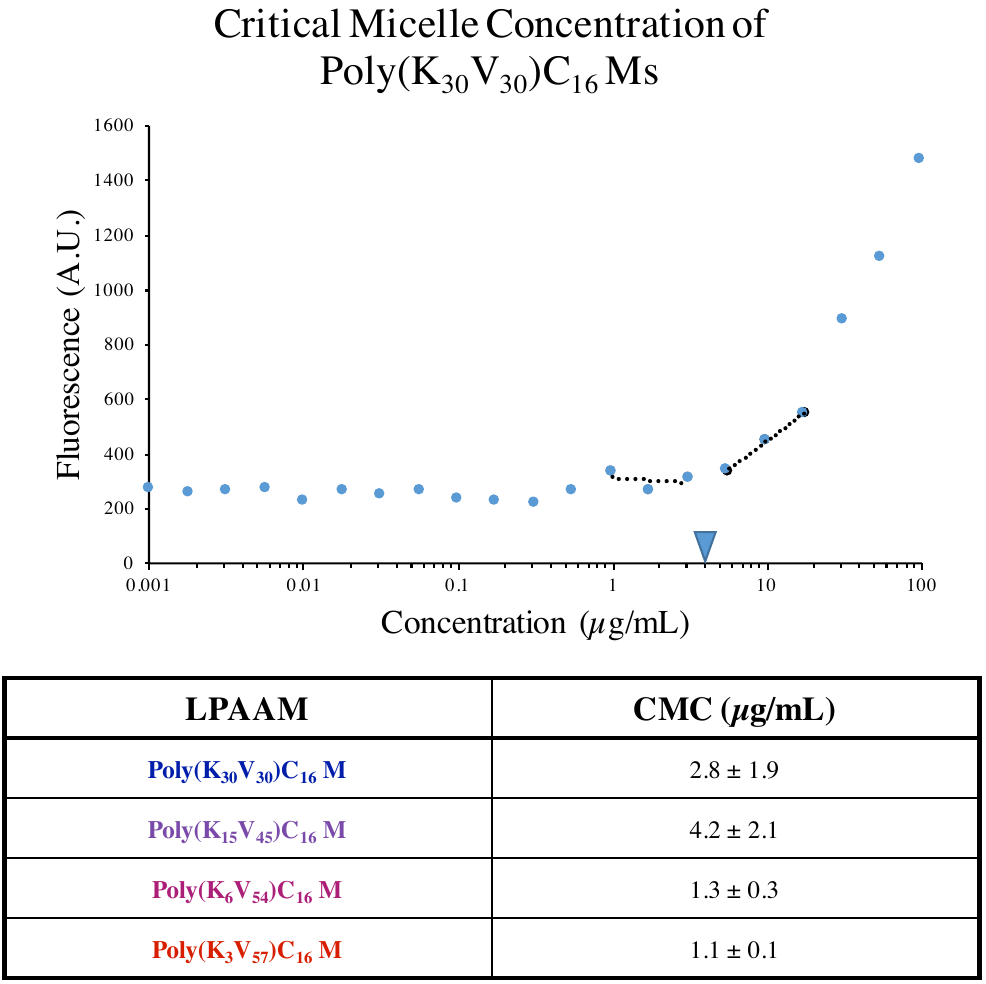

**Figure S8:** LPAA self-assembly characterization. The critical micelle concentration (CMC) of each formulation was evaluated by serial dilution of a 100 μg/mL solution of LPAA in PBS with 1 μM 1,6-diphenyl-1,3,5-hexatriene (DPH). The DPH fluorescence inflection point is considered the minimum concentration at which micelles have spontaneously formed. All formulations had similar CMCs which were consistent batch-to-batch. The tabulated values given are the averages from three independent CMC studies for each LPAA.

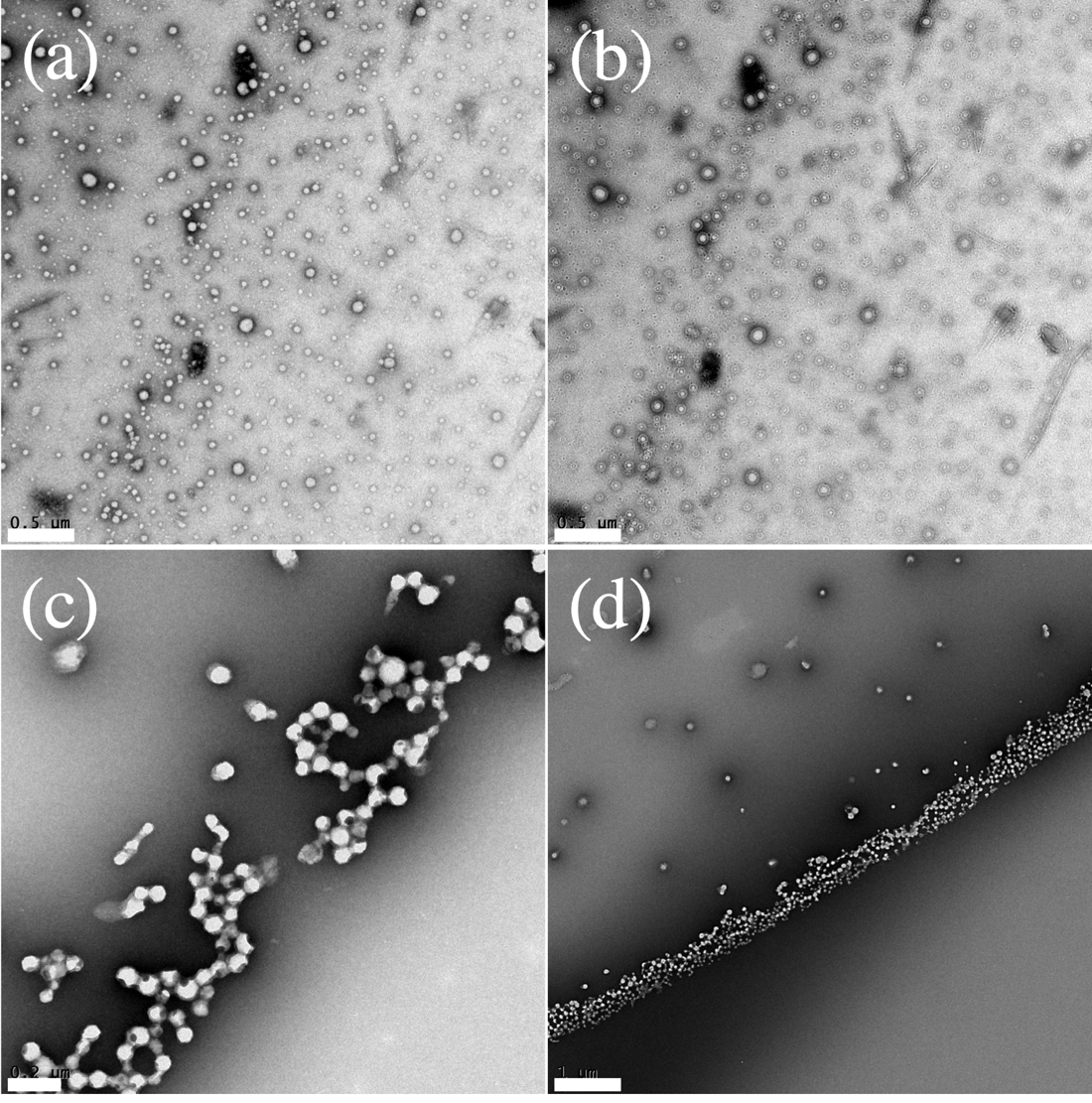

**Figure S9:** LPAAM TEM micrographs for Poly(K_6_V_54_)C_16_ and Poly(K_3_V_57_)C_16_. (a) Solubilized Poly(K_6_V_54_)C_16_ was found to form small nanoparticles with a double-wall as shown in the (b) out-of-plane micrograph. (c) - (d) Poly(K_3_V_57_)C_16_ was found to form larger, solid core nanoparticles. Scale bars are (a) 0.5 µm, (b) 0.5 µm, (c) 0.2 µm, and (d) 1 µm.

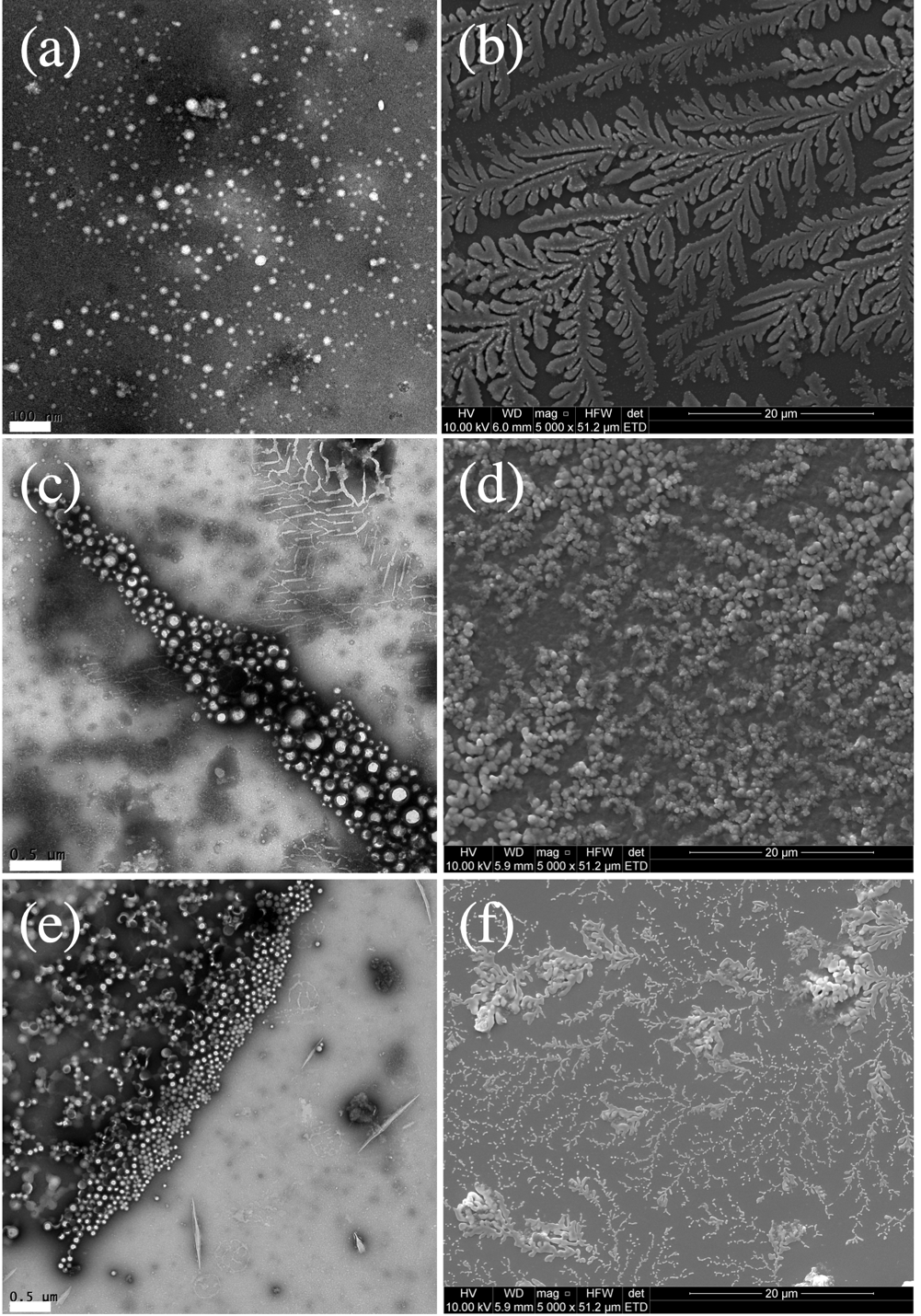

**Figure S10:** LPAAM EM micrographs. (a), (c), (e) TEM and (b), (d), (f) SEM micrographs for (a) and (b) Poly(K_30_V_30_)C_16_, (c) and (d) Poly(K_15_V_45_)C_16_, and (e) and (f) Poly(K_6_V_54_)C_16_ are shown. SEM and TEM analysis of the three nanoprecipitate formulations showed interesting differences. Notably, all samples on SEM contained branching higher-order structure that remained regardless of the sample processing method employed (*data not shown*). TEM was chosen as the preferred method for evaluation as nanoparticles were easily distinguished from each other. TEM micrograph scale bars are (a) 0.1 µm, (c) 0.5 µm, and (e) 0.5 µm. (b), (d), and (f) SEM micrograph scale bars are each 20 µm.

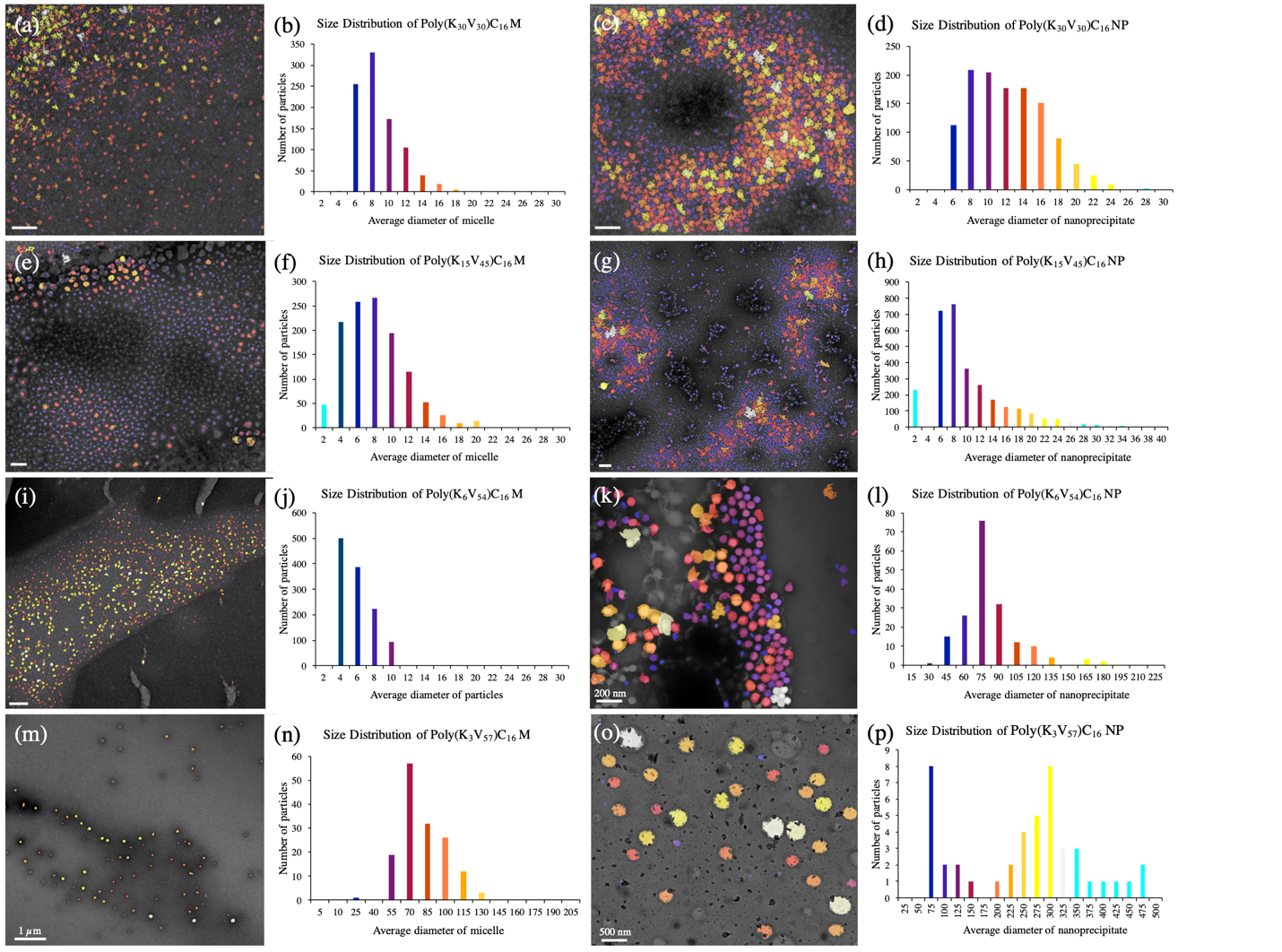

**Figure S11:** LPAAM and LPAANP image-based automated size analysis. TEM micrographs were analyzed using the Bruker ESPRIT 2.0 software package for which micrograph contrast and brightness as well as expected nanoparticle circularity and diameter were adjusted for each analysis. Every run was directly compared with the original micrograph to ensure the accuracy of automated sizing. Micrographs overlaid with ((a), (c), (e), (g), (i), (k), (m) & (o)) the analysis color scheme and ((b), (d), (f), (h), (j), (l), (n) & (p)) particle size distributions are shown. Each plot is labeled with its corresponding formulation with the matched micrograph to its left. For example, (a) shows the micrograph overlay for poly(K_30_V_30_)C_16_ Ms with (b) its size distribution and (c) displays the micrograph overlay for poly(K_30_V_30_)C_16_ NPs with (d) its corresponding distribution. Micrograph scale bars are 50 nm except where noted *(i.e*. (k), (m), and (o)). The size distribution x-axis varies by plot where necessary to show the overall nanoparticle distribution profile.

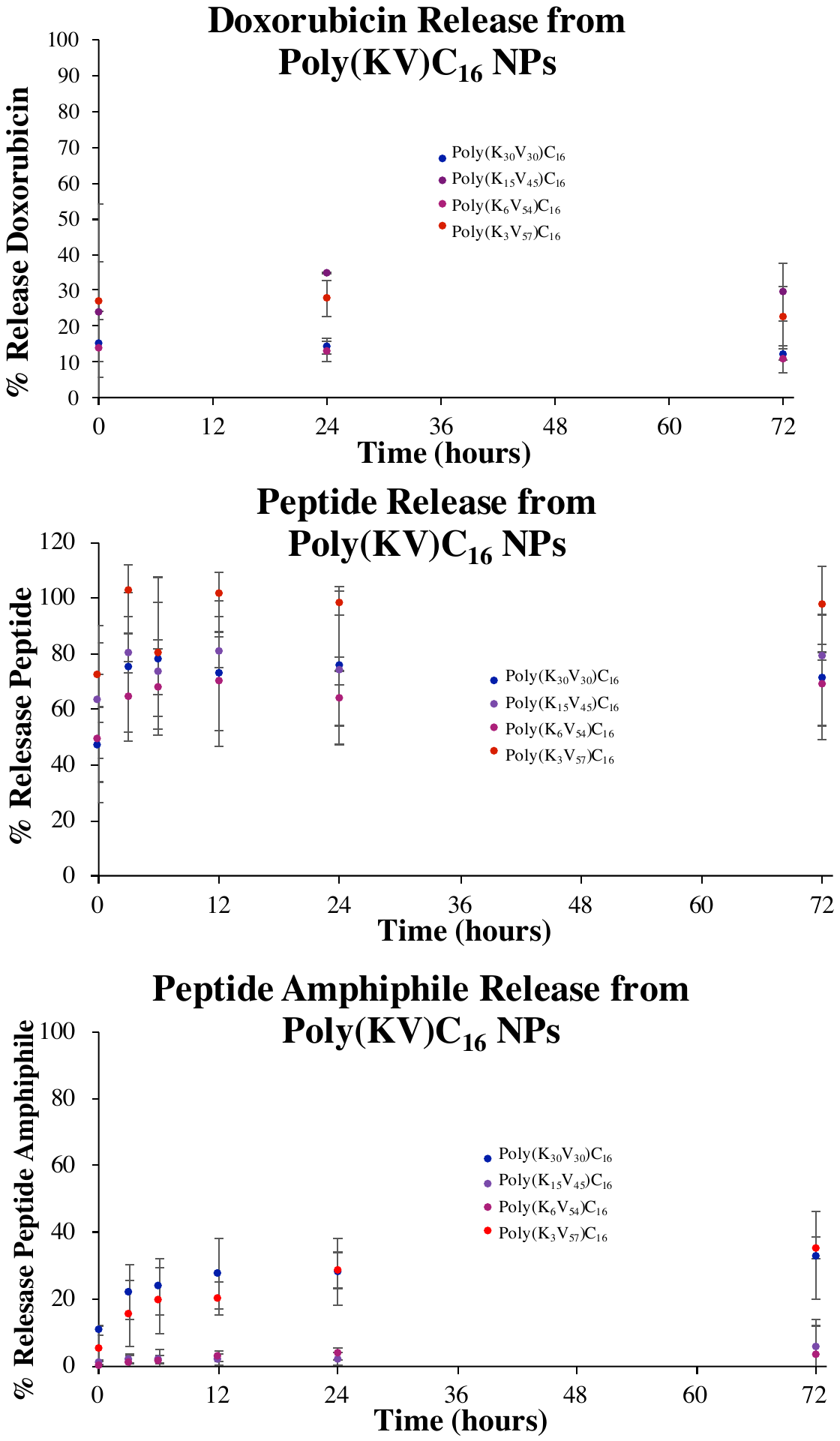

**Figure S12:** LPAANP associated drug release profiles. Payload release was evaluated by incubation in PBS and subsequent molecular weight cutoff filtration. Some burst release (*14%*  -*100%*) was observed for all drug associated nanoprecipitate formulations except PA-loaded Poly(K_15_V_45_)C_16_ NPs and Poly(K_6_V_54_)C_16_ NPs. Controlled gradual drug release over time was only seen with PA-loaded Poly(K_30_V_30_)C_16_ NPs and Poly(K_3_V_57_)C_16_ NPs.

**Figure S13:** Maleimide conjugation analysis. Poly(K_6_V_54_)C_16_ Dox NPs are hereafter referred to as LPAANP_6/54_ Dox for simplicity. FTIR spectra reveal characteristic peak changes after LPAANP_6/54_ malemide surface modification and further reaction with thiolated antitail DNA. First, a shift in the broad N-H peak from ~ 3170 cm^-1^ (purple) to 3000 cm^-1^ (orange and grey) indicates a change from primary to secondary amine indicating successful attachment of maleimide-DEG groups to nanostructure surface primary amines. Additionally, the appearance of peaks at 1650 cm^-1^ (C=C), 1450 cm^-1^ (C=O), 1080 cm^-1^ (N-C), and 835 cm^-1^ (H-C-C-H) indicate the presence of the maleimide group.

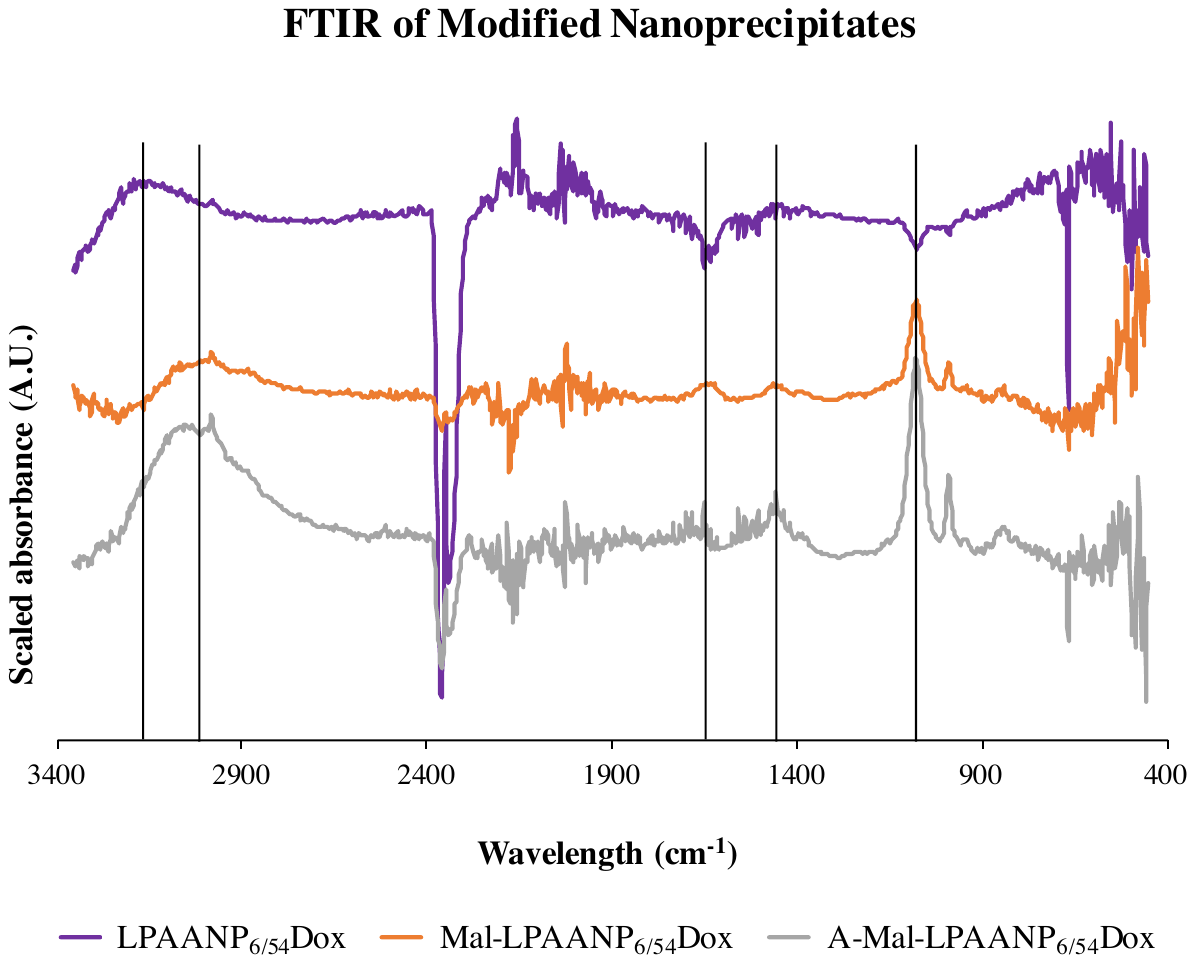

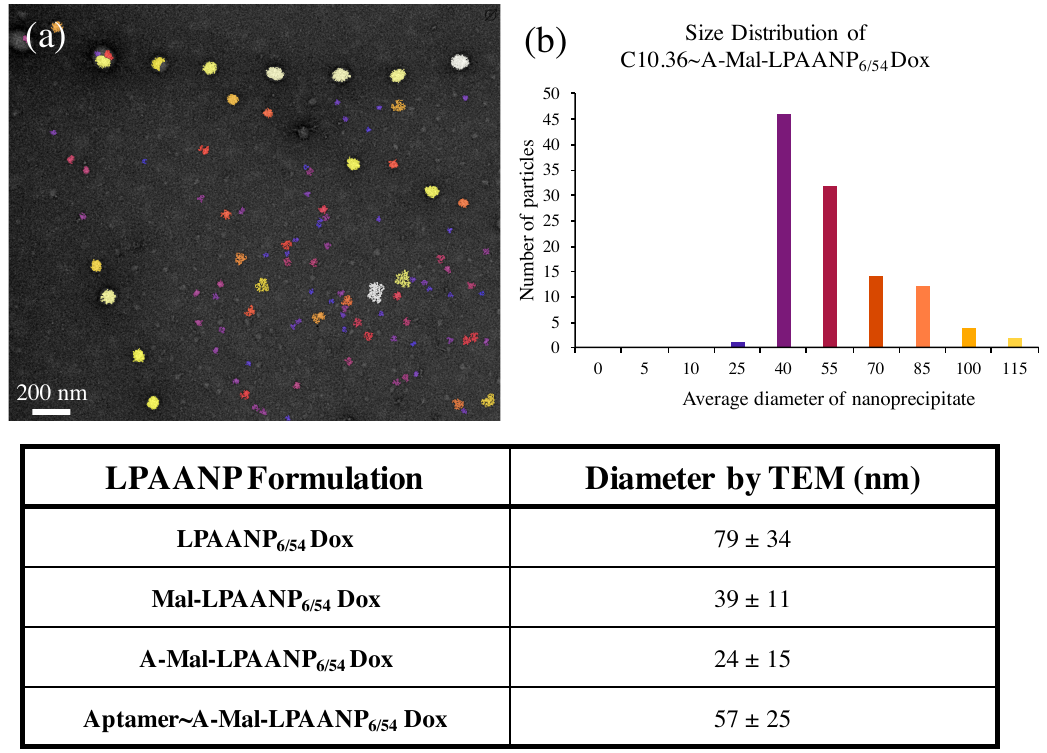

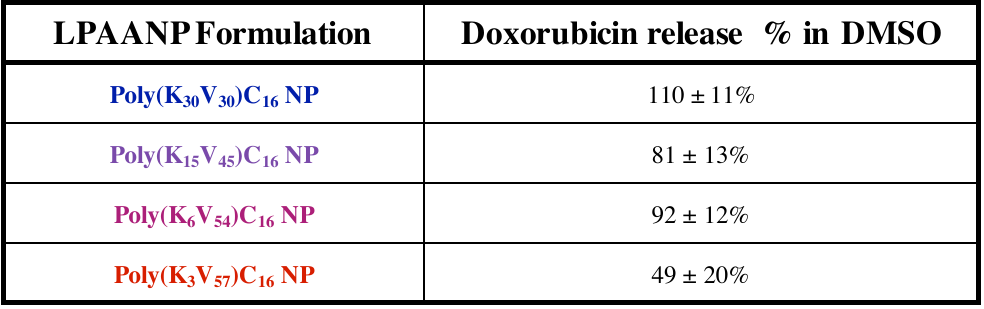

**Figure S14:** LPAANP_6/54_ Dox image-based automated size analysis. Using the previously described Bruker ESPRIT 2.0 software package, (a) a TEM image was analyzed to determine (b) the nanoprecipitate size of LPAANP_6/54_ Dox. This was repeated for (c) progressively surface modified nanoprecipitates through aptamer conjugation for which no significant morphological nor size differences were observed. Each formulation particle size was generated from three TEM micrographs taken of independently produced samples.

**Table S2:** LPAANP Dox release efficiency. Total release efficiency was evaluated by diluting nanoprecipitate in 97.5% dimethylsulfoxide (DMSO) / 2.5% phosphate buffered saline (PBS) to a final concentration of 10 µg/mL. The concentration of each entrapped species was calculated from standards of each material in the same diluting solution. Dox was found to have reasonable release efficiencies regardless of formulation chemistry.

**Table S1:** Aptamer and antitail DNA sequences. All sequences shown are listed 5' to 3'.

| **C10.36** | CTAACCCCGGGTGTGGTGGGTGGGCAGGGGGGTTAGCGACGACGACGACGACGACGA |
| --- | --- |
| **G24A** | CTAACCCCGGGTGTGGTGGGTGGACAGGGGGGTTAGCGACGACGACGACGACGACGA |
| **scApt** | GCCATTGCCATTGCCATTGCCATTGCCATTGCCATTGCCATTGCCATTGCCATTGCGACGACGACGACGACGACGA |
| **antitail** | TCGTCGTCGTCGTCGTCGTCG |
